# Non-sequential alignment of binding sites for fast peptide screening

**DOI:** 10.1101/2023.08.01.551496

**Authors:** Frédéric Guyon, Gautier Moroy

## Abstract

**Motivation:** Peptides are molecules involved in many essential biological activities by interacting with proteins in the body. They can also be used as therapeutic molecules, particularly to disrupt protein-protein interactions involved in disease. However, it is difficult to identify potential therapeutic peptides if the structure of the protein-protein complex to be inhibited has not been solved.

**Results:** The PepIT program was developed to propose peptides that can interact with a given protein. PepIT is based on a non-sequential alignment algorithm to identify peptide binding sites that share geometrical and physicochemical properties with a surface region of the target protein. PepIT compares the entire surface of the target protein with the peptide binding sites of the Propedia dataset, which contains more than 19,000 high-resolution protein-peptide structures. Once a peptide binding site similar to a portion of the protein surface is found, the peptide bound to the binding site is repositioned on the corresponding portion of the protein surface.

**Availability and implementation:** The PepIT source code is freely available at https://github.com/DSIMB/pepit

**Supplementary information:** Supplementary data are available online.

## 1. Introduction

Peptides are historically defined as a polypeptide chain consisting of 2 to 50 amino acids. Peptide-protein interactions are essential for diverse biological events, such as growth regulation, stress response, and immune signaling in all living organisms. They are estimated to represent up to 40% of the Protein-Protein Interactions (PPIs) within the cell [1]. Recently, there has been increased interest in the design of peptide-based drugs that bind and inhibit target proteins involved in pathologies [2, 3].

To this purpose, effective computational methods to identify peptides capable of binding to a given protein and to understand what are the structural mechanisms responsible for the interaction between a peptide and a protein are required.

To identify the molecular mechanism responsible for a protein-peptide interaction, protein-peptide docking methods are often used. These methods can be roughly divided into three approaches [4]. The template-based docking (TBD) method is based on the similarities, mainly the sequence identity, of experimental structures to model the structures of protein-peptide interactions. The TBD methods have low computational cost that enables to use it for large-scale applications. The success of TBD methods depends strongly on the existence of known structures, whose numbers are fortunately constantly increasing year by year. The second approach is local docking, which focuses on a site of interest on the protein surface. The methods developed to perform local docking of peptides require the user to define the site of interest with the highest possible accuracy based on experimental or computational data, or both. However, it is still computationally expensive to perform local docking with a large library of peptides that vary in sequence and size. In addition, if no data on the target site is available, multiple binding sites must be considered, further increasing the computational cost. Finally, it is possible to perform docking calculations on the entire surface of the protein, this approach is called global docking. Although approximations can be used to represent the protein or peptide to reduce computational time, this approach is inherently more computationally intensive than the previous ones. In general, peptides are very flexible, which is one of the reasons why peptide docking is so difficult. Current peptide docking approaches have difficulty to predict correctly the conformation of peptides in the vicinity of the target protein, especially if they are larger than ten residues.

Recently AlphaFold [5] has been used to predict the structure of multimers [6] and in particular the structure of protein-peptide complexes [7, 8, 9]. Docking and AlphaFold-based methods require knowledge of the sequence of the peptide whose interaction with a protein is to be studied. There are few approaches capable of designing a *de novo* peptide based only on the knowledge of the structure of a target protein. The method EvoBind designs peptides from target and peptide sequences. It is based on AlphaFold to model the complex structure and performs an evolution of the peptide sequence directed by the scoring of AF.

The approach described in this paper, named PepIT, identifies peptides that are likely to interact with a given protein target. Depending on the chosen protein target, these peptides can be protein activity regulators, protein-protein interaction inhibitors, antimicrobial peptides, or peptides acting as antibodies with specific binding characteristics and high affinity for targeted proteins. It has been established that two similar binding sites can interact with similar small molecules [10]. Thus, by searching for local similarities of the target protein surface with a collection of complexes, it is possible to identify peptides candidates for bounding interaction. PepiT selects these peptides from structures available in the PDB as natural peptides or protein fragments. It makes use of a structure comparison algorithm that allows for rapid screening of binding site data banks.

The binding sites comparison is based on a sequence order-independent alignments that are obtained by searching for cliques in product graphs. Clique algorithms have been used in CavBase [11], eF-site [12], SuMo [13], PocketMatch [14], SOIPPA [15], SiteEngine [16], eMatchSite [17] and ProBis [18]. Most of the above approaches compare or align binding sites only and require prior identification of the location of binding sites on the surface of the protein. Less numerous approaches (SOIPPA, ProBis) can compare whole protein and detect local similarities on complete protein’s surfaces, but they only align alpha carbon atoms.

PepiT searches for local similarities with known binding sites on the entire surface of the target protein. The PepiT procedure used to search for structural similarities differs from the above mentioned methods based on a clique approach. It does not require the prior knowledge of the location of the binding sites. For this goal, it makes use of a clique algorithm [19] combined to a maximal weight matching search. The binding sites have variable conformation and this flexibility makes geometrical comparison difficult. Moreover, all-atom comparisons of proteins counting a few thousand atoms lead to some very large product graphs and algorithms for computing cliques, such as the Bron-Kerbosch algorithm, run in exponential time for general graphs. Clique algorithms are intractable even for moderate size binding sites searches on whole protein surface considering all atoms. They also aim to maximize clique size and not a similarity score. To tackle these difficulties, mappings between atoms are weighted. The weights allow for a higher precision on alpha carbons with a greater tolerance for other atoms, especially for side chain atoms. PepiT computes non sequential alignments of a subset of atoms (*α* carbons or *α* or backbone atoms) with a clique approach, and modifies and extends these alignments by searching for maximal weight matching in a bipartite correspondence graph. Hence, the PepiT comparison method is not too sensitive to the side chain atom’s position and is able to account for target flexibility.

The target protein surface is compared to a bank of reference binding sites. The 3D alignment allows the superimposition of the retrieved binding site and therefore, since the binding site is associated with a bound peptide, it proposes a pose of this peptide to the unbound protein structure.

The assessment of the method is performed on three benchmarks of proteinpeptide complexes: PeptiDB [20], PepPro [21] and Propedia [22]. The first two data sets are used to adjust the parameters of the algorithm and to estimate the probability distribution of the Pepit score, the last one is a large collection of 19,000 peptide-protein complexes used to show the efficiency of Pepit on the target protein BCL-XL as a test case.

sectionData

### 1.1. PeptiDB

PeptiDB is a non-redundant data set containing 108 high quality structures of peptide-protein complexes [20]. The peptide sequence lengths is ranged from 5 to 15 residues. The structures were resolved by X-ray crystallography, with a resolution better than 2.0Å. 47 unbound protein structures are available. PeptiDB is the reference data set for assessing bioinformatics tools devoted to the study of the protein-peptide interactions.

### 1.2. PepPro

More recently, PepPro, a database of experimentally determined peptide-protein complexes, has been released [21]. It contains 89 non-redundant peptide-protein complex structures. The peptides have a size between 5 and 30 residues. For 58 peptide-protein complexes, the unbound protein structures are provided.

### 1.3. Propedia

Propedia is a database of protein-peptide complexes containing over 19,000 high-resolution structures [22]. The peptide sizes ranged from 2 to 50 amino acids. The structures of the peptide binding sites with the associated peptides were downloaded for comparison with protein surfaces. If sufficient structural and physicochemical homology is detected between a peptide binding site from Propedia and part of the surface of a given protein, we can assume that the peptide is able to bind to this part of the surface. PepiT then places the peptide on the corresponding site of the studied protein so that it can be studied further by the user.

## 2. Methods

### 2.1. The binding site banks

The algorithm implemented in PepIt tries to find the best matching between binding site atoms *S*_1_ and target surface atoms *S*_2_ by maximising the clique edge-weight.

A target protein is given with no known interacting peptide or for which peptides are known, but no complex with this protein exists in the PDB. It is also be possible to predict new interactions located elsewhere than on a known binding site or, for a given binding site, to obtain different peptides able to bind to the protein. For these purpose, we build binding site libraries from different set of protein-peptide complexes from PeptiDB (103 non redundant complexes) [20] PepPro (89 complexes) and Propedia (19,000 high-resolution structures).

For each complex, the receptor binding site is extracted by retaining the atoms located at a cutoff distance from the peptide. To study the effect of the cutoff, different sets are constructed with the distance varying from 4Å to 10Å by steps of 1Å. We thus obtain 7 banks. Moreover, when an atom is selected, the C*α* of the residue to which this atom belongs is added to the binding site.

### 2.2. The PepiT matching algorithm

PepiT computes a non-sequential alignment, that is, it proposes a pairing of atoms that come from residues that are not necessarily in the same order in the two structures. The alignment minimizes the distortion between the two sets of aligned atoms. The distortion is defined here as the sum of the differences between all the internal distances of the structures. Hence, the alignment aims at preserving the internal distances within a cutoff threshold that controls the flexibility allowed in the alignment. The best alignments are selected and following these alignments, the binding sites found in the bank are superposed with the targeted protein surface. The superposition performed by a translation and a rotation. The same rotation and translation are also applied to the peptide associated with the binding site, which is thus positioned on the surface of the protein.

The described algorithm hereunder aims to compute matching between two sets of typed atoms *S*_1_ and *S*_2_. All atoms are typed with respect to their physicochemical properties. A link or a match between two atoms exists only if they are of the same type. In this case, the atoms are said to be compatible. We consider all pairs of compatible atoms *i* = (*i*_1_*, i*_2_) and *j* = (*j*_1_*, j*_2_), the first one from *S*_1_ and the second one from *S*_2_. The distortion between atoms *i*_1_ and *j*_1_ of the structure *S*_1_ and atoms *i*_2_ and *j*_2_ of the structure *S*_2_:

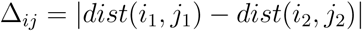

where *dist*(*i*_1_*, j*_1_) is the euclidean distance between atoms *i*_1_ and *j*_1_. A matching between *S*_1_ and *S*_2_ denoted by **M** consists in a set of atom pairs (*i*_1_*, i*_2_), *i*_1_ *∈ S*_1_ and *i*_2_ *∈ S*_2_, where **M** gives a one-to-one mapping between *S*_1_ and *S*_2_.

The mean distortion is:

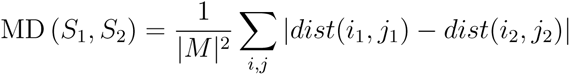

PepiT algorithms aims to compute the largest matching such that the mean distortion between structures is below a given threshold noted Δ*_max_*.

The parameter Δ*_max_* allows to control the amount of flexibility. We can be interested in a long alignment with a moderate mean MD, more specifically where all distortions are not too large, or in a shorter alignment where all distortions are below a stringent threshold. It should be noticed that, in contrast to the RMSD calculation, the MD distance can be computed without any prior optimal superposition. The MD needs internal distances computations that do not depend on the location and the orientation of the proteins.

Problem 2 is solved in two steps. First, a maximum clique approach is applied to the correspondence graph. Nodes of the correspondence graph are pairs *i* = (*i*_1_*, i*_2_) of atoms associating an atom of *S*_1_ to an atom of *S*_2_ and edges connect nodes *i* to nodes *j* if *i*_1_! = *i*_2_ and *j*_1_! = *j*_2_ and the distortion Δ*_ij_* between *i* and *j* is less than a Δ*_max_* fixed to 1Å. It should noted that this graph is not defined as the usual product graph of two protein graphs, which is usually the basis of clique approach. The correspondence graph is constructed with a subset of atoms to limit the computation time. This subset consists in *C_α_* atoms only or in backbone atoms. In order to have enough atoms to perform the calculations, we systematically add the alpha carbons of the residues whose atoms participate in the binding sites. Maximum cliques are searched in this graph with a Bron-Kerbosh algorithm [19]. Cliques are then filtered to eliminate mirror similarities. Clique gives a subset of matching atoms in both protein structures with approximately conserved inter-atomic distances. One of the two structural elements can be the mirror image of the other. Indeed, a structure and its mirror-symmetric image have identical inter-atomic distances. Such a mirror symmetry can be detected by a simple matrix determinant calculation. After removal of mirror similarities, remaining cliques with at least four nodes in common are merged.

The clique approach yields a solution where all the distortions have an upper bound of Δ*_max_*. The *C_α_* are supposed to undergo limited movements between bound and unbound conformations. Therefore, similarities between binding sites are more easily detected with the *C_α_*alone without the side atoms that can undergo larger displacements under binding.

Second, we consider the bipartite graph with edges connecting compatible atoms of *S*_1_ to atoms of *S*_2_.

Edges of the bipartite graph are weighted as follows:

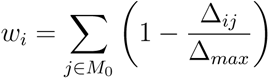

A matching is a set of edges where no two edges have a common vertex.

The total weight of a matching *M* is

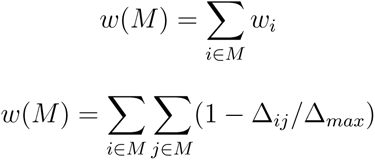

where *n* is the number of atoms in the alignment that is *n* = *|M |*. We have

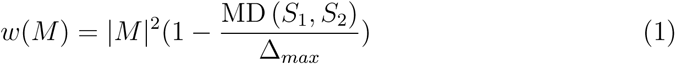

For the final best matching, we always have:

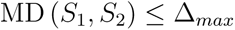

Efficient algorithms exist to get matching of maximum size or of maximum constant weight. A push-relabel algorithm can be used to find a maximal matching with constant weights. The main difficulty of problem 2 is that each edge weight depends on the matching *M* . To our knowledge, no algorithm exists that solves this problem exactly. We propose an alternate iterative algorithm, which we prove to converge to a local maximum (cf. Suppl). Given a clique *M*_0_ computed in the precedent step, a matching that maximizes the score MDscore (*S*_1_*, S*_2_) is performed.

It can be shown that a matching between *S*_1_ and *S*_2_ maximizing *w*(*M*) is a solution of the following optimization problem:

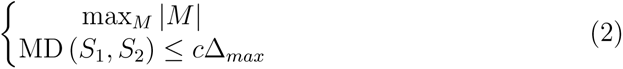

with a given constant *c*,0 *< c ≤* 1

#### Algorithm 1 Maximum weight matching

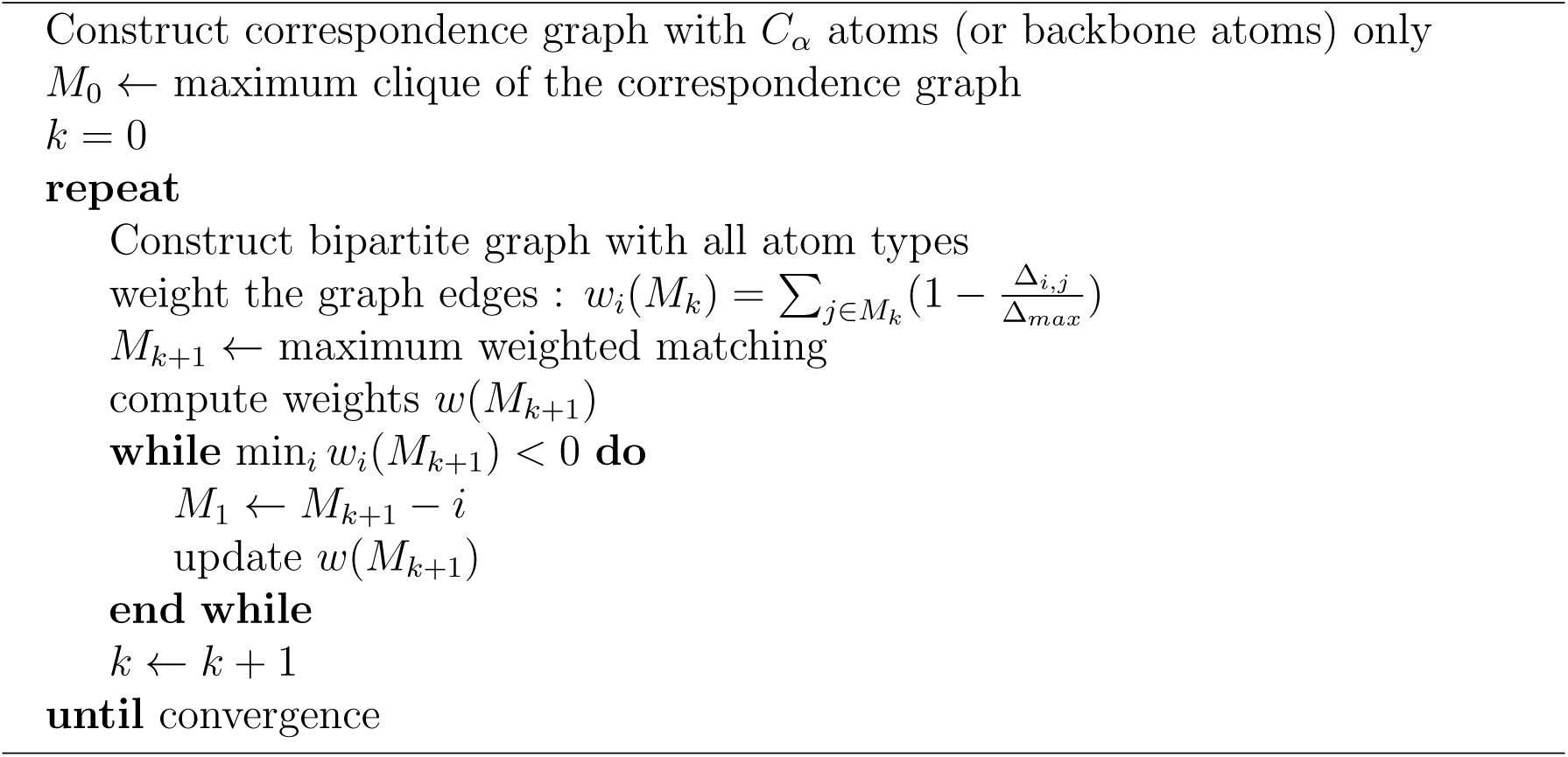

### 2.3. Tuning parameters and score analysis with PeptiDB and PepPro

A difficulty arises when using the matching weight as a ranking score (equation 1). The matching weight value depends on the size of the binding site and therefore on the size of the bound peptide. Thus the score produced by a small peptide that can bind to the target could be lower than a score obtained with a longer peptide but bound to a site that has weak or no similarity with the target. It implies that the mapping score implies a poorer discrimination for small binding sites (see Figure 1).

**Figure 1:**
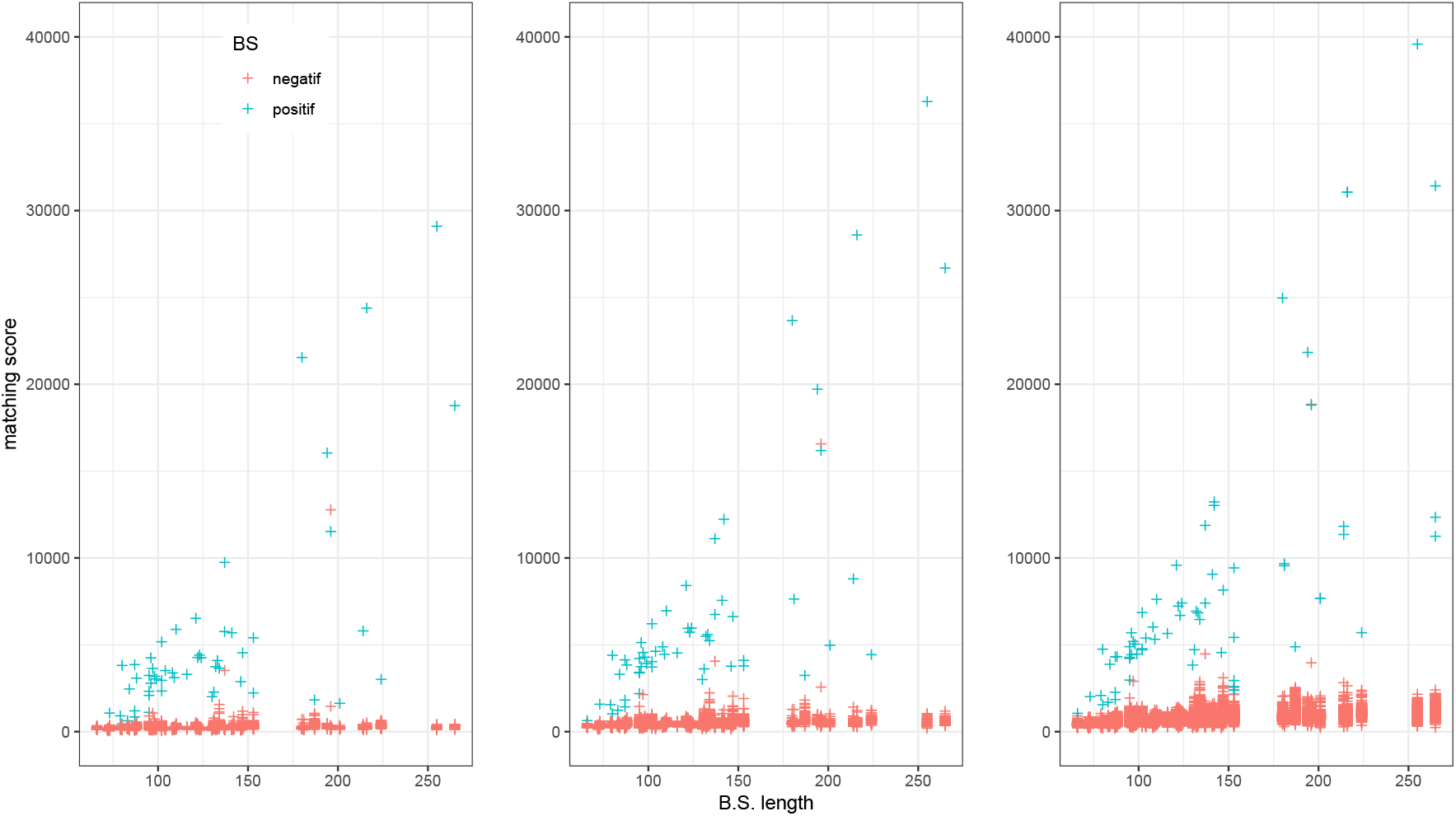
Matching score values versus binding site lengths for different precision values. From left to right Δ*_max_* = 1, 1.5, 2. Negative binding sites are represented by red points and positive ones by blue points

First the score’s dependence on the size of the binding sites is analyzed on the search results obtained on two banks of protein-peptide complexes PeptiDB and PepPro. Both benchmarks consist of a non redundant representative reduced set of protein-peptide complexes. As such, they allow the adjustment of parameters suitable for more extensive binding site searches. This allows to choose a scoring with a distribution independent of the binding size.

Second, the algorithm parameters are adjusted to optimize the recognition of binding sites of the two banks. The adjustments concern the addition of alpha carbons, the contact distance threshold and the precision parameter Δ*_max_*. We recall that the algorithm searches in a library of binding sites for the most similar to a surface patch of the targeted protein. The peptides associated with these binding sites are then candidate peptides. To assess the algorithm efficiency in finding effective peptides, we use two subset of the complex bank consisting of proteins bound to a peptide for which an apo form also exists. For a given apo form, the binding site of the corresponding complexed protein is considered positive. All others are tagged as negative binding sites. The test consists in comparing all apo forms against all binding sites. It is successful when the true binding site of the holo form is ranked first with the highest score.

It is a difficult task for proteins that undergo major conformational changes when bound to a peptide. Another difficulty for the comparison algorithm is due to the many equivalent arrangements of atoms and residues that exist in proteins, especially on protein surfaces, which leads to incorrect matches. It is sufficient that a few identical residues contribute to the binding sites to obtain such false matches.

## 3. Results

### 3.1. Alignement length distribution

The algorithm is a heuristic to approximately solve the optimization problem (2) which aims to maximize the length of the alignment for which the average distortion is below a given threshold Δ*_max_*. For positive binding sites, the length of alignments grows with site size. In contrast, the size of alignments with negative binding sites increases much less relatively to site size and to threshold Δ*_max_* (Figure 2). This length on average depends linearly on the precision parameter Δ*_max_*. Dividing the alignment length by Δ*_max_* normalizes the alignment length and makes its distribution almost independent of the Δ*_max_* parameter (Figure 3) of the binding site size (Figure 4). The normalized length criterion enables positive binding sites to be selected with high specificity. The highest specificity is obtained for small Δ*_max_*.

**Figure 2:**
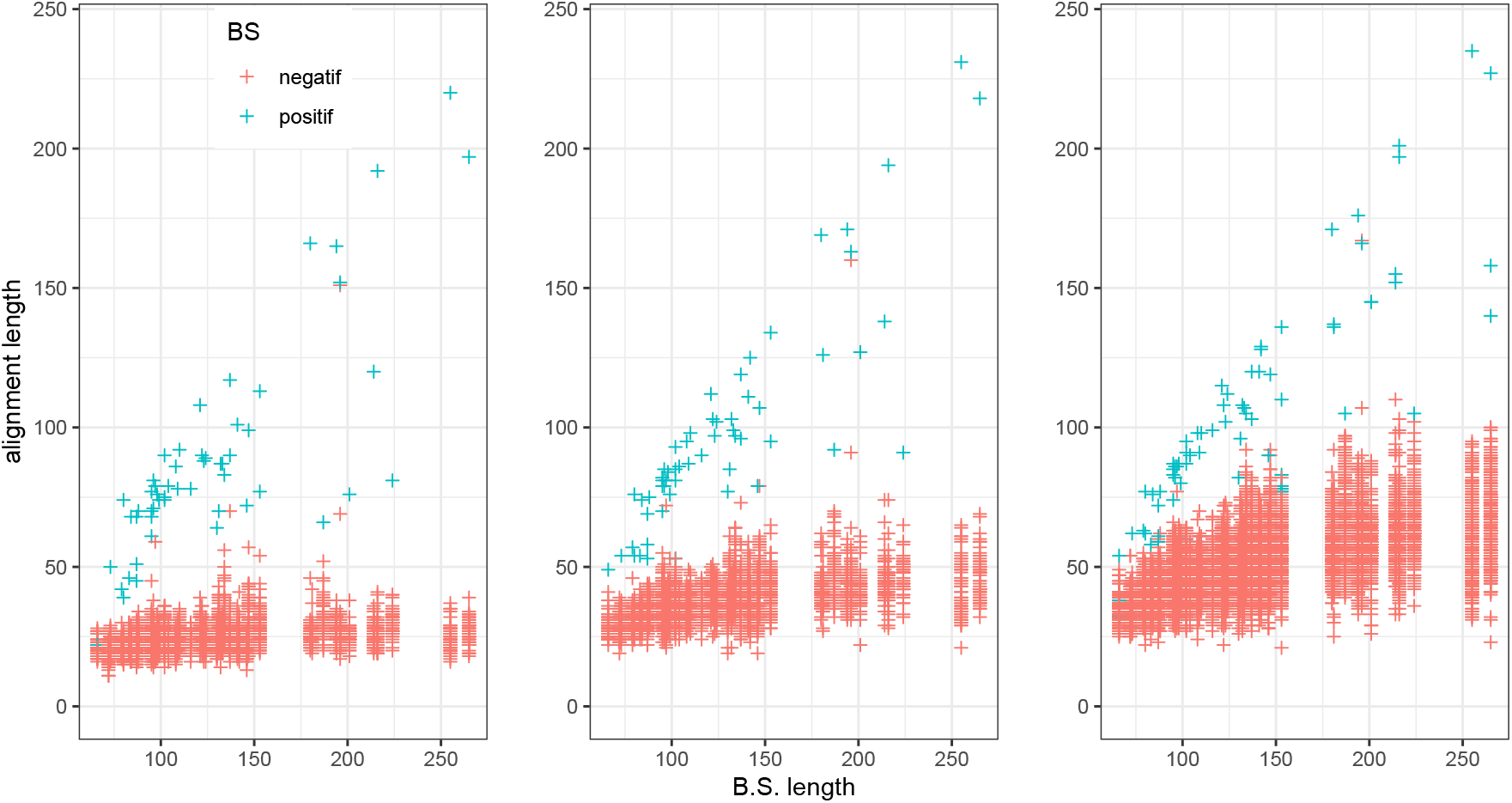
Alignment lengths versus binding site length for different precision values. From left to right Δ*_max_* = 1, 1.5, 2. Negative binding sites are represented by red points and positive ones by blue points

**Figure 3:**
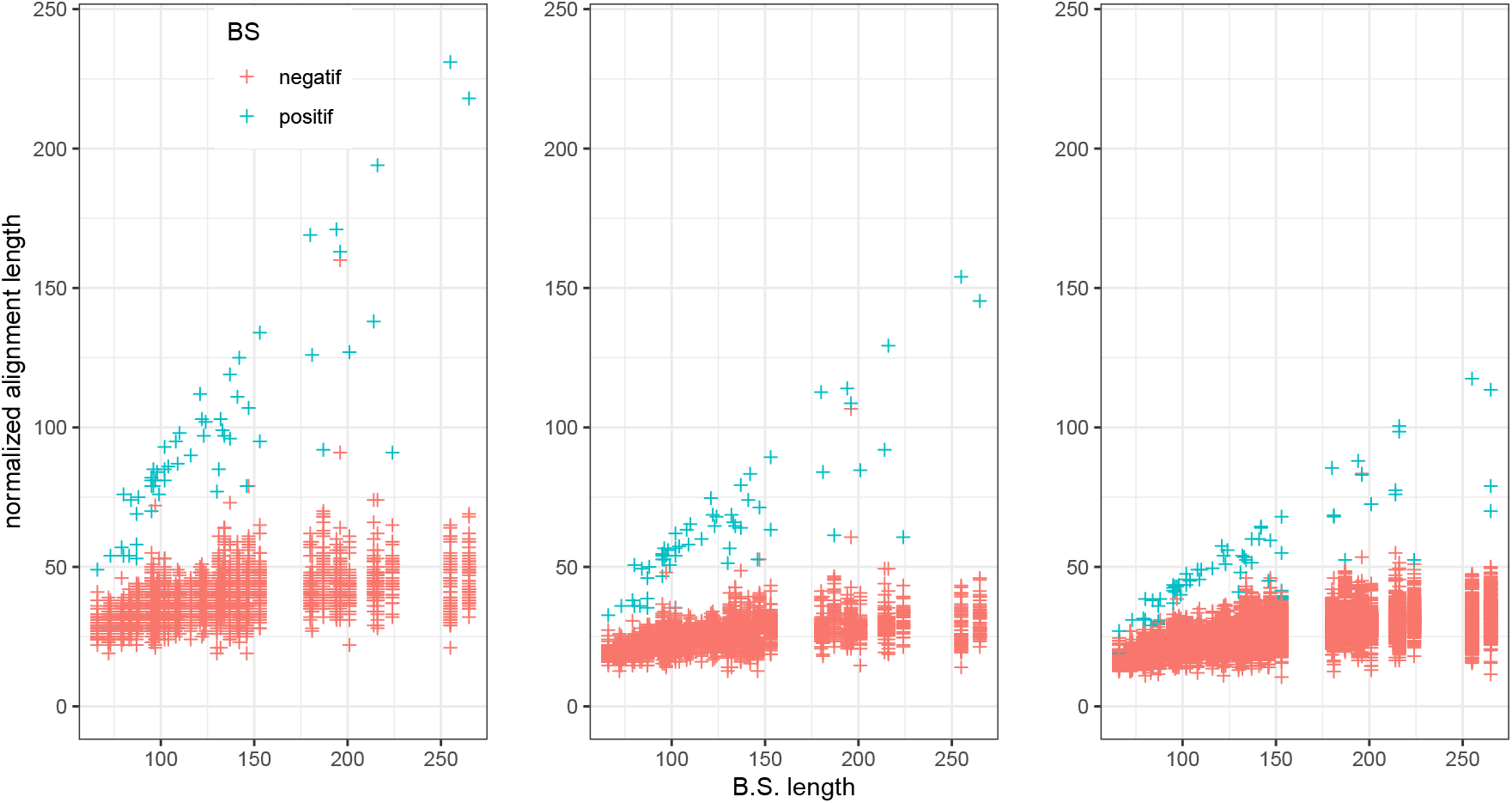
Normalized alignment lengths versus binding site length for different precision values. From left to right Δ*_max_* = 1, 1.5, 2. Negative binding sites are represented by red points and positive ones by blue points

**Figure 4:**
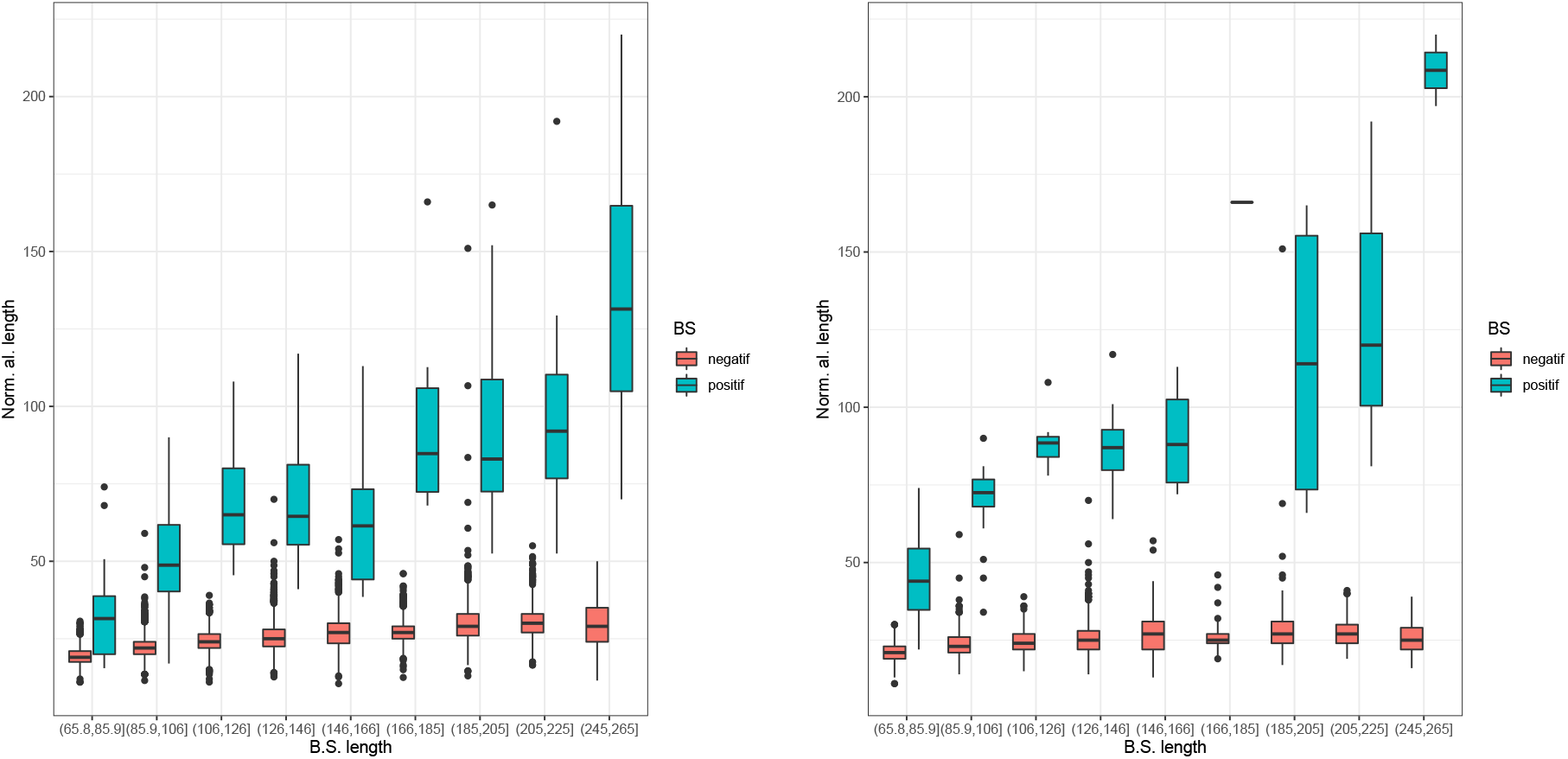
Boxplots representing from left to right, distributions of normalized alignment lengths for increasing range of binding site lengthes.

**Figure 5:**
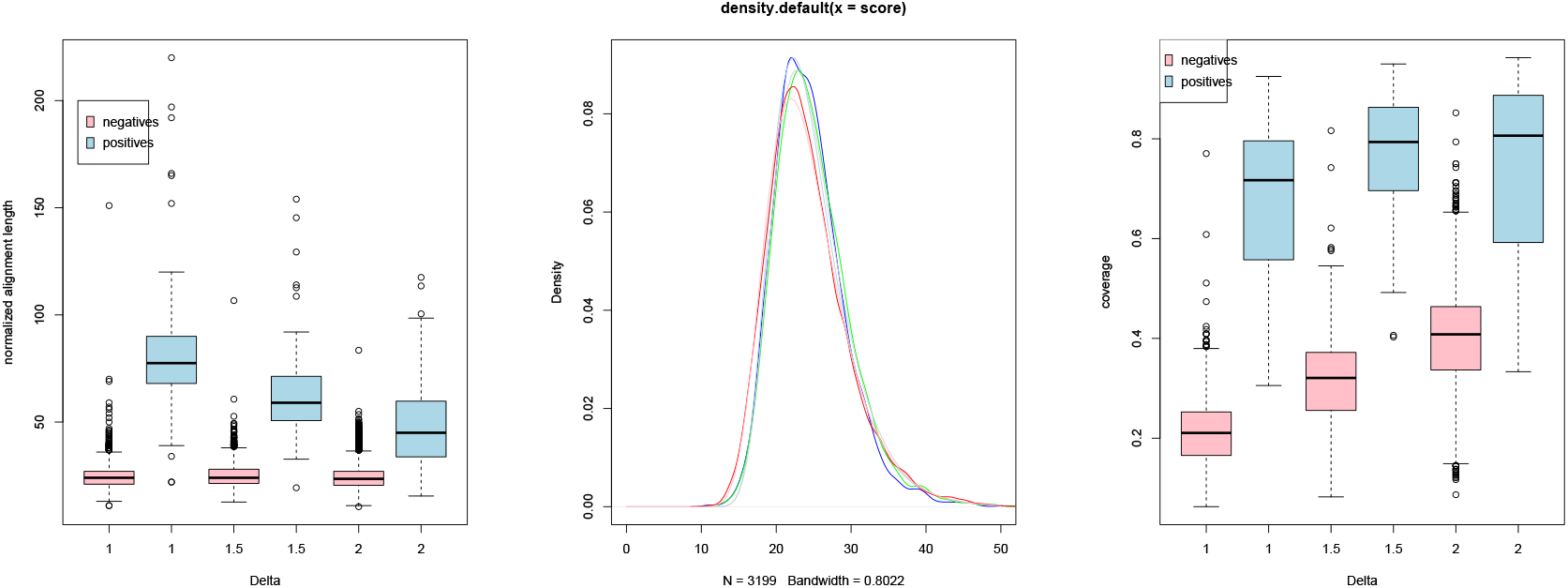
Boxplots representing from left to right, distributions of normalized alignment lengths, Gumbel distributions and distributions of coverage for Δ*_max_* = 1, 1.5, 2.

The Fisher-Tippett theorem states that the maximum of a sequence of independent and identically distributed variables, under certain conditions of convergence, follows a distribution belonging to one of the three possible laws: Gumbel’s law, Fréchet’s law or Weibull’s law. These laws belong to the class of generalized extremum laws [23]. For a given Δ*_max_*, the length of negative alignments is the maximum value obtained for each binding site on a protein target. The set of scores obtained during a comparison of the surface of a target protein against the library of binding sites, allows the estimation of the parameters of the distribution. These distributions are characterized by three parameters: the location *µ*, the scale *σ* and the shape parameter *ξ*. We first estimated these parameters on the binding sites given by the PeptiDB complex library. Parameter estimation is obtained with the extRemes function library which allows fitting a generalizes extreme value distribution to an empirical distribution of score values [24]. Table 1 shows that that the shape parameter *ξ* is close to zero and therefore the distribution of the maximum alignment length approaches a Gumbel distribution. A simple estimate of the parameters of the Gumbel distribution is given by the following calculations : For a sequence of normalized alignment lengths from negative binding sites *l_i_*, let *m* be the average score and *s* the stan<u>d</u>ard deviation of the scores. The scale and the location can be estimated by 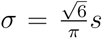 and *µ* = *m − σγ*, where *γ* = *−*0.57721566 *. . .* is the Euler’s constant. Therefore, the probability that a normalized alignment length *l* is greater than the observed length is given by 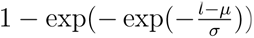.

**Table 1:** Generalized value distribution parameters fit on the normalized alignment length for different Δ*_max_* values.

| $\Delta_{max}$ | loc | scale | shape |
| --- | --- | --- | --- |
| 1.0 | 22.32 | 4.020 | 0.0081 |
| 1.5 | 22.64 | 4.13 | -0.010 |
| 2 | 21.86 | 4.42 | -0.02 |

### 3.2. Coverage distribution

Figures 6 show that the distribution of the coverage of the positive binding sites varies due to the small number of positive samples compared to the negative ones, but does not significatively augment with the binding site. The coverage tends to decrease with the negative ones. The reason is that, as for negative binding sites the alignment length is approximately constant for different binding site sizes, necessarily the coverage decreases with the binding site size. Thus the gap between the alignment lengths of positive and negative sites grows with the binding site length. This implies that it becomes easier to recognize good binding sites when their size increases.

**Figure 6:**
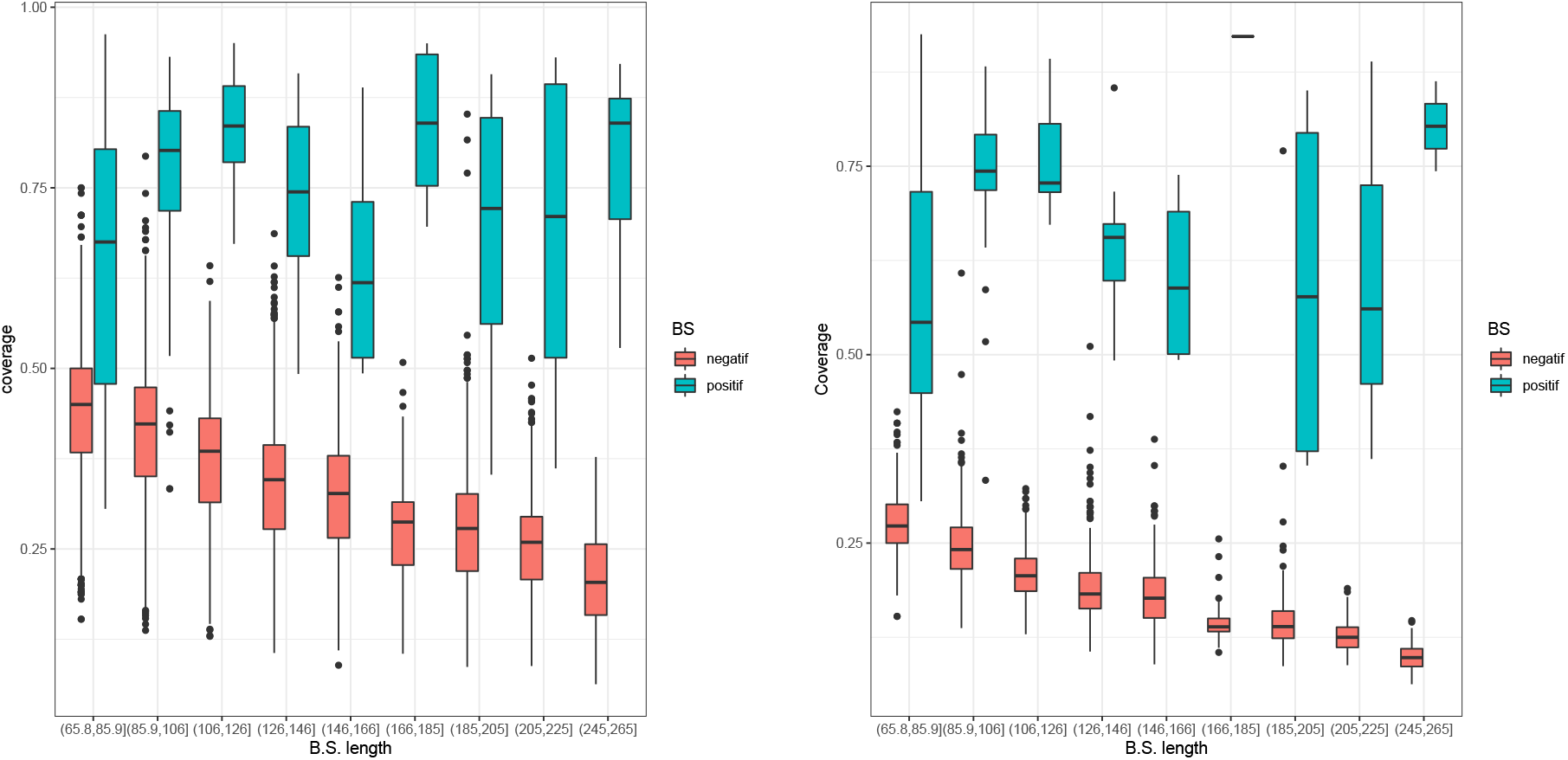
Boxplots representing from left to right, distributions of coverage for Δ*_max_* = 1*Å* and of all coverages with Δ*_max_* = 1, 1.5, 2, 2.5*Å*.

**Figure 8:**
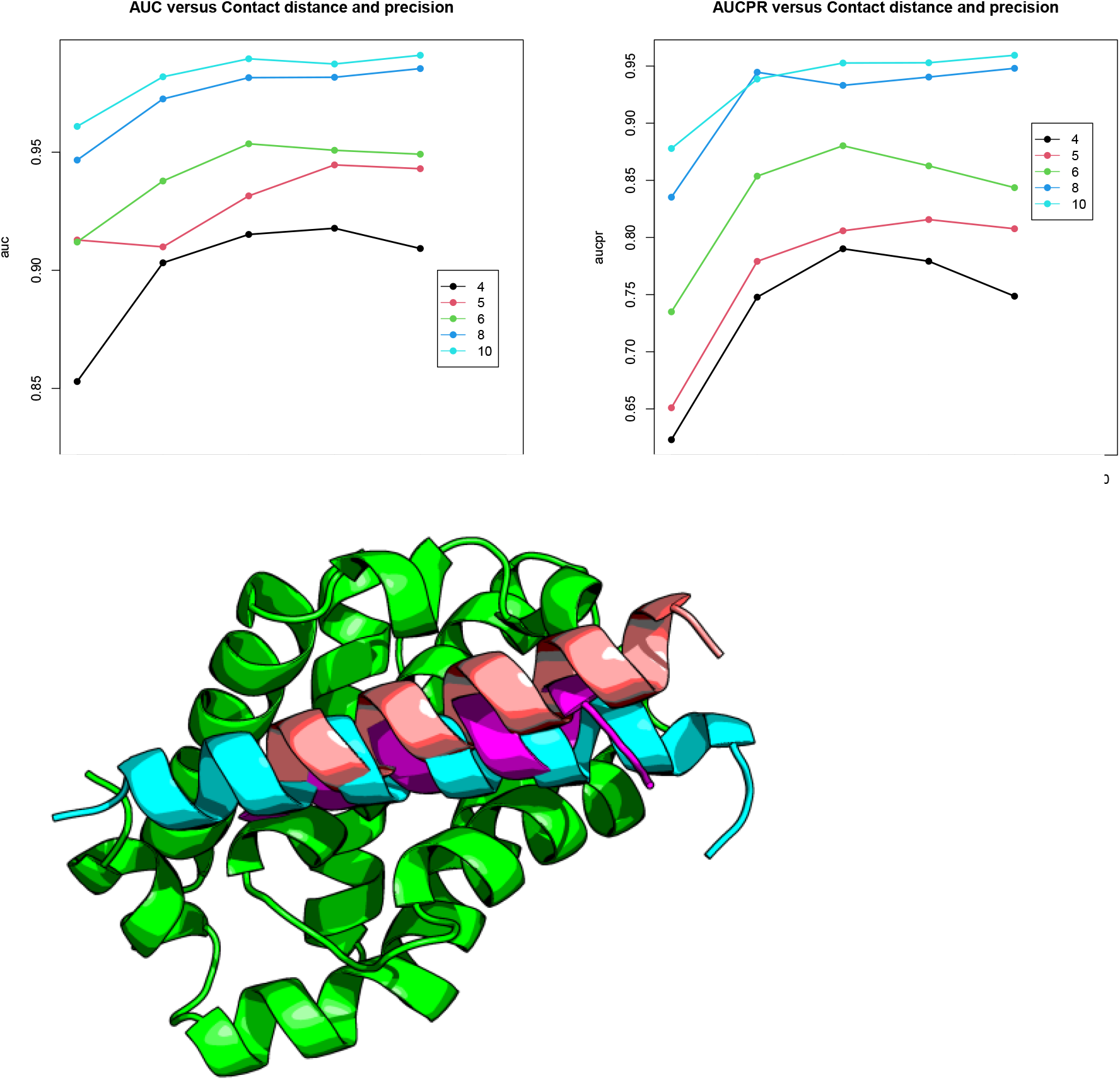
Cartoon representation of Bcl-xL, in green, in interaction with BH3 peptide from BAD in cyan (PDB ID:1G5J). The CAMKII*α* peptide identified and positioned on Bcl-xL by PepIT is shown in magenta. The CAMKII*α* peptide in red results from the structural prediction of the complex Bcl-xL / CAMKII*α* peptide by the AlphaFold-Multimer program.

By definition of the optimization problem 2, the length of the alignment, and therefore the coverage, automatically increases with higher precision parameter values. Negative binding sites are more sensitive to an increase in Δ*_max_* than positive sites, whose coverage varies much more moderately. The ratio of spurious matches get higher with Δ*_max_* but is limited in positive binding sites. Therefore, the coverage criterion enables positive binding sites to be selected with greater sensitivity, regardless of their length.

The coverage is not the result of a maximization procedure, it does not follow an extreme value distribution but Figure 6 shows that it is independent of the precision parameter and of the binding site length.

In conclusion, a P-value is computed from the Gumbel distribution of the normalized alignment length. This P-value is used to discard the potentially negative solutions and the coverage is used to classify the remaining solutions and hence to select the best putative positive binding sites.

### 3.3. Tuning of the parameters

The algorithm parameters are adjusted using ROC and precision-recall analysis to improve the recognition of binding sites. A ROC curve plots the true positive rate (or sensitivity) versus the false negative rate (1-specificity) for all possible score values. The area under this curve called AUC is used to assess the effectiveness of the parameters. The precision-recall curves plots the true positive rate versus the predicted positive value that is the fraction of true positive binding sites among the retrieved ones. The area under this curve is called AUCPR. The adjusted parameters of the search are the binding site interaction distance *d_contact_* varying from 4 to 10Å, wether or not C*α* atoms are added to the binding sites and the precision tolerance parameter Δ*_max_*. The corresponding apo and holo proteins are the same proteins or close homologous proteins. Thus, the set of matching atoms is known, and it is possible to evaluate the precision of the search. This benchmark also assesses the ability of the method to find the correct location of the binding site on the surface of the apo protein. The AUC and AUCPR are shown in tables 2 and 3 and displayed in figure 7. When the contact distance is increased, binding sites contains a greater number of atoms and also a greater number of atoms not involved in the interaction with the peptide. Their position can be more variable and it is more difficult to align them with the equivalent atoms of the unbound apo protein and the alignment coverage decreases. When this threshold is released, i.e. when the threshold Δ*_max_* increases, automatically the length of the matching increases but beyond a certain value, the algorithm matches atoms which are not in the true alignment. In case the protein does not undergo large deformations in the presence of the peptide, it is close to the binding site size.

**Table 2:** AUC values for different Δ*_contact_* values and for binding sites with additional *α*-carbons (+)

| Distorsion | 0.5 | 1 | 1.5 | 2 | 2.5 |
| --- | --- | --- | --- | --- | --- |
| Contact | AUC |  |  |  |  |
| 4 | 0.895 | 0.869 | 0.851 | 0.876 | 0.889 |
| 4+ | 0.853 | 0.903 | 0.915 | 0.918 | 0.909 |
| 5 | 0.892 | 0.845 | 0.856 | 0.865 | 0.873 |
| 5+ | 0.913 | 0.910 | 0.931 | 0.945 | 0.943 |
| 6 | 0.899 | 0.828 | 0.842 | 0.831 | 0.835 |
| 6+ | 0.912 | 0.938 | 0.954 | 0.951 | 0.949 |
| 8 | 0.959 | 0.886 | 0.896 | 0.899 | 0.898 |
| 8+ | 0.947 | 0.973 | 0.982 | 0.982 | 0.985 |
| 10 | 0.902 | 0.910 | 0.957 | 0.940 | 0.935 |
| 10+ | 0.961 | 0.982 | 0.990 | 0.987 | 0.991 |

**Table 3:**
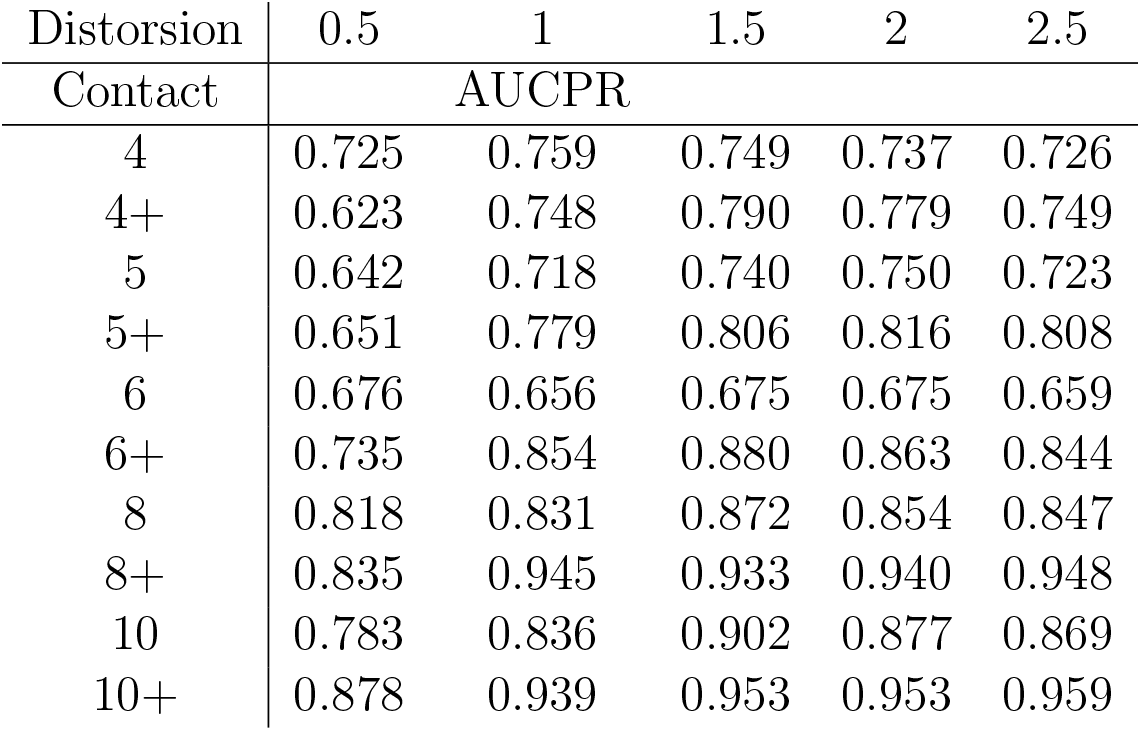
AUCPR for different *Delta_contact_*values and for binding sites with additional *α*-carbons (+)

**Table 4:** Peptide able to interact with Bcl-xL according to PepIT calculations.

| Peptide | PDB ID | Peptide chain | mean distortion | al. length | coverage | P-value |
| --- | --- | --- | --- | --- | --- | --- |
| BAD | 2BZW | B | 0.0000 | 153 | 1.0000 | 1.10e-13 |
| SOUL | 3R85 | H | 0.5416 | 95 | 0.6376 | 4.46e-08 |
| Bcl2-L-2 | 4CIM | Q | 0.4987 | 36 | 0.6207 | 2.22e-02 |
| HBx | 5B1Z | C | 0.5486 | 84 | 0.5753 | 5.16e-07 |
| Beclin-1 | 5VAU | F | 0.4658 | 81 | 0.5625 | 1.01e-06 |
| PUMA | 2YJ1 | B | 0.4673 | 91 | 0.5515 | 1.09e-07 |
| BID | 4QVE | B | 0.5046 | 94 | 0.5497 | 5.57e-08 |
| BIM | 5C3G | B | 0.4423 | 78 | 0.5493 | 1.96e-06 |
| Bcl2-L-11 | 4A1U | B | 0.4412 | 88 | 0.5301 | 2.12e-07 |
| Bak | 5FMJ | B | 0.4824 | 91 | 0.5291 | 1.09e-07 |
| TCTP | 4Z9V | D | 0.5224 | 78 | 0.5000 | 1.96e-06 |
| DHDH | 5UA5 | B | 0.5715 | 57 | 0.4524 | 2.10e-04 |
| BAX | 4BD6 | C | 0.7123 | 54 | 0.3857 | 4.09e-04 |
| BMF | 5TZQ | C | 0.5926 | 50 | 0.3731 | 9.96e-04 |
| egl-1 | 1TY4 | D | 0.5534 | 62 | 0.3246 | 6.90e-05 |
| BAK-2 | 5TWA | C | 0.5012 | 50 | 0.3226 | 9.96e-04 |
| dF4 | 6MBC | B | 0.6731 | 44 | 0.2667 | 3.78e-03 |
| CAMKIIα | 1CM1 | B | 0.5462 | 31 | 0.1384 | 6.61e-02 |

The addition of alpha carbons to the binding sites greatly improves the recognition with significantly increased AUC and AUCPR. The reason is that alpha carbons are less likely to move during binding with a peptide and are therefore more easily found by the PepiT algorithm. When alpha carbons are present in the binding site, increasing the contact distance also produces an improvement in aucpr. The recognition is therefore significantly improved with this strategy of including alpha carbons at the binding sites. Alignments resulting from cliques computed with additional C-*α* carbons are more accurate and the clique extension phase is able to include a larger set of well matched atoms. Increasing the contact distance increases the size of the binding sites to be compared and increasing the precision parameter increases the size of the graphs produced. Thus, increasing the values of these two parameters improves the recognition but implies a strong increase of the computation time. It is therefore necessary to find a compromise between recognition efficiency and computation time. This improvement is weak for precision parameter values higher than 1 Åand for contact distances higher than 8. We have therefore chosen the following parameters for the algorithm: addition of the alpha carbons, a contact distance of 8Å (or 10) and a precision of 1Å(or 1.5).

The two libraries used in this benchmark are composed of proteins with nonredundant sequences. However, some retrieved binding sites that are not part of the holo form of the library show remarkable similarities with the target protein. Nevertheless, they are not classified as positive binding sites.

## 4. Case study: Bcl-xl

Bcl-xL is a protein belonging to the Bcl2 family of proteins This protein family is well-known to participate to the apoptosis regulation. Bcl-xL exerts its antiapoptotic role by interacting with pro-apoptotic proteins, such as Bax or Bak. Bcl-xL is composed by four BH (Bcl-2 Homology) domains. The Bcl-xL general structure is characterized by nine *α*-helices, with a hydrophobic *α*-helix core surrounded by amphipathic helices. Overexpression of the Bcl-xL protein is known to induce chemoresistance to cancer therapies. The disruption of the interaction between Bcl-xL and pro-apoptotic proteins by inhibitors can restore the apoptosis in cancer cells. The development of molecules specifically targeting Bcl-xL is therefore a promising approach to cancer therapy. To propose new interfering peptides, we chose the structure of Bcl-xL, which forms a complex with the BH3 peptide derived from the BAD protein (PDB ID:2BZW [25]). The BAD peptide was deleted before searching in the Propedia database to determine whether peptides may interact with Bcl-xL. Among the peptides best classified by PepIT, most were in the *α*-helical structure. The top-ranked peptides are derived from anti-apoptotic proteins of the Bcl-2 family, which are known to biologically interact with Bcl-xL or other anti-apoptotic proteins of the Bcl-2 family. Moreover, these peptides are inserted into the groove corresponding to the site that interacts with pro-apoptotic proteins, such as BAD, Bak, Beclin-1 or Bim. PepIT is therefore able to recognize peptides derived from partners that can interact with Bcl-xL, while positioning them correctly on its surface.

Interestingly, PepIT computations showed that the Bcl-xL binding site has structural similarities with calmodulin (PDB ID: 1CM1[26]), a protein implicated in many crucial biological processes in eukaryotic cells, such as inflammation, apoptosis, and immune response. The sequence similarity between calmodulin and Bcl-xL is approximately 44.0%. To check if these proteins have a similar structure, the TM-score of 0.29 indicates that these proteins do not share the same fold. Therefore, identifying a structural similarity between the two peptide binding sites on these protein is difficult based on sequence or structure alignments alone.

Based on these structural similarities between the peptide binding sites, PepIT identified that a peptide of 18 residues derived from calcium/calmodulin-dependent protein kinase type II alpha (CAMKII*α*) might be able to bind to the Bcl-xL binding site. The sequence similarity between CAMKII*α* peptide and BAD BH3 peptide is 42.0%. The CAMKII*α* peptide has a *α*-helical structure similar to that of BH3 peptides. Interestingly, CAMKII*α* is known experimentally to interact with BCL2-associated athanogene 5 (BAG5) and BCL2-associated athanogene 6 (BAG6) proteins [27].

To confirm that the CAMKII*α* peptide might be able to bind Bcl-xl, AlphaFold-Multimer was used to model the interaction between the CAMKII*α* peptide and Bcl-xl. Only the CAMKII*α* peptide and the Bcl-xl sequences were provided. The average pLDDT score [28] for the best model proposed by AlphaFold-Multimer is of 81.87, indicating that the confidence in this model is high. Although no data on the location of the binding site was provided, AlphaFold-Multimer modeled the peptide in *α*-helical structure and interacted at the same location on the surface of Bcl-xl as the BAD peptide (Fig. 3).

Therefore, the hypothesis that the CAMKII*α* peptide interacts with Bcl-xl is fully supported by the AlphaFold Multimer results. This peptide seems then to be able to compete with the BH3 peptides and, consequently, with pro-apoptotic proteins. It can be assumed that this peptide is a good candidate for the restoration of apoptosis in tumor cells.

## 5. Conclusion

PepiT uses only the structure of the protein to compare its surface with known protein-peptide interaction sites from the ProPedia database. Peptides may target a protein without specifying a site on its surface. Pepit score is independent of the peptide size so that small peptides can be proposed. Furthermore, our results show that PepiT is tolerant to conformational changes of the interaction sites due to the intrinsic flexibility of the proteins or to the adaptation of the position of the residues to the presence of a ligand. Thus, PepiT can be utilized to discover a peptide that is capable of interacting with a particular protein of interest. Subsequently, the identified peptides can be enhanced using other specialized programs to improve its affinity [29].

The application of PepIT to the identification of peptides capable of interacting with Bcl-xl allowed the highlighting of a peptide derived from CAMKII which has a secondary structure identical to that of the peptides known to bind to Bcl-xl. Interestingly, the CAMKII peptide shares a weak sequence identity with peptides known to bind Bcl-xl and the calmodulin, the protein with which it interacts, also shares a weak sequence identity with Bcl-xl. PepIT has therefore succeeded in finding a new peptide that could potentially bind to Bcl-xl. This result is supported by the fact that AF2 predicts with high confidence a structure in which the interaction between the CAMKII peptide and Bcl-xl takes place at the site on the surface of Bcl-xl known to bind peptides.

The main limitation of PepIT is related to the limited diversity of peptides that have been experimentally solved and are available in the ProPedia database or more generally in the PDB data bank. However, with the intensive use of new methods based on artificial intelligence, which are able to predict the structure of proteins and complexes with great accuracy, the number of peptides considered will no longer be a problem, if it is possible to access the results of these predictions.

---

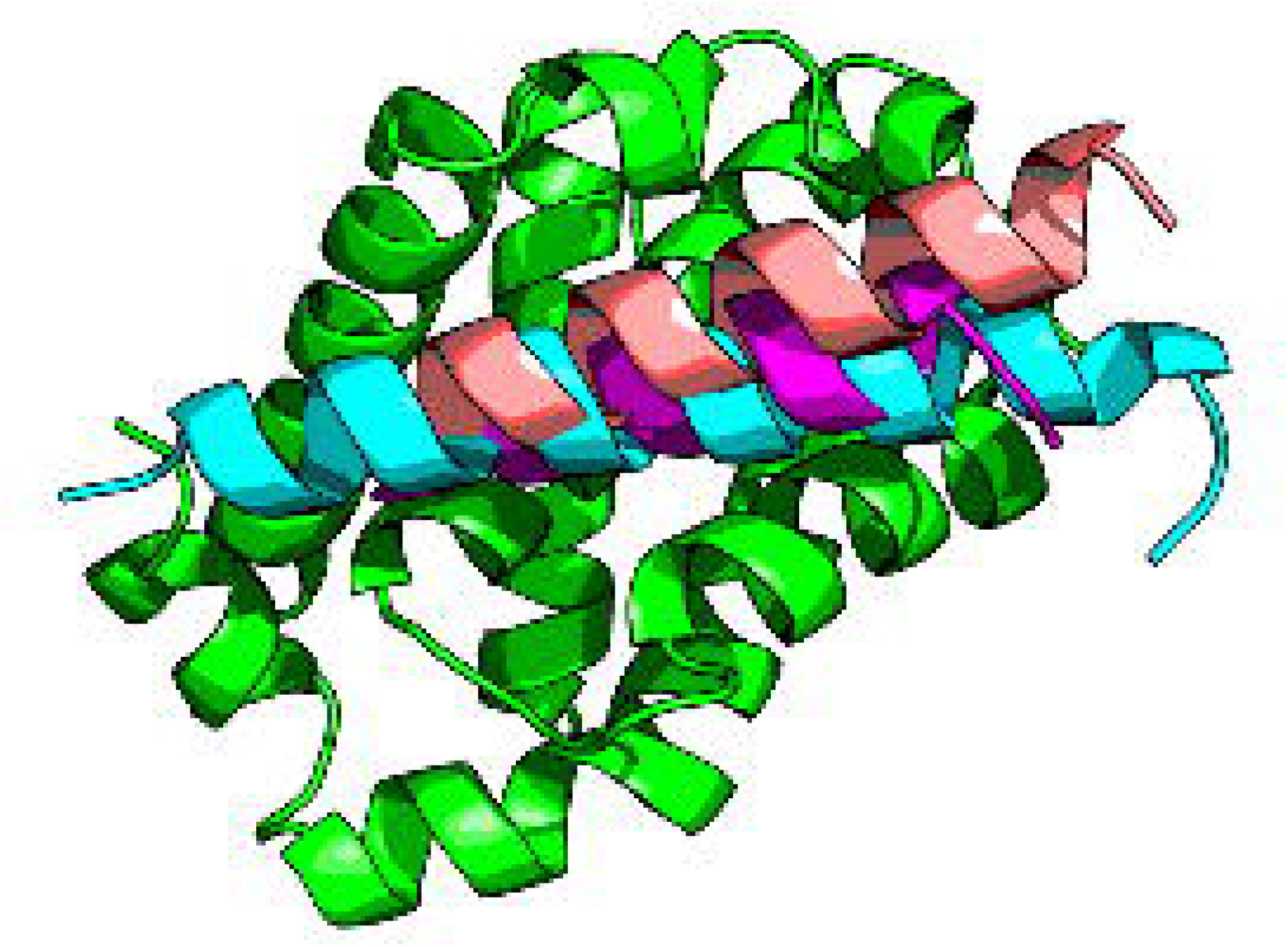

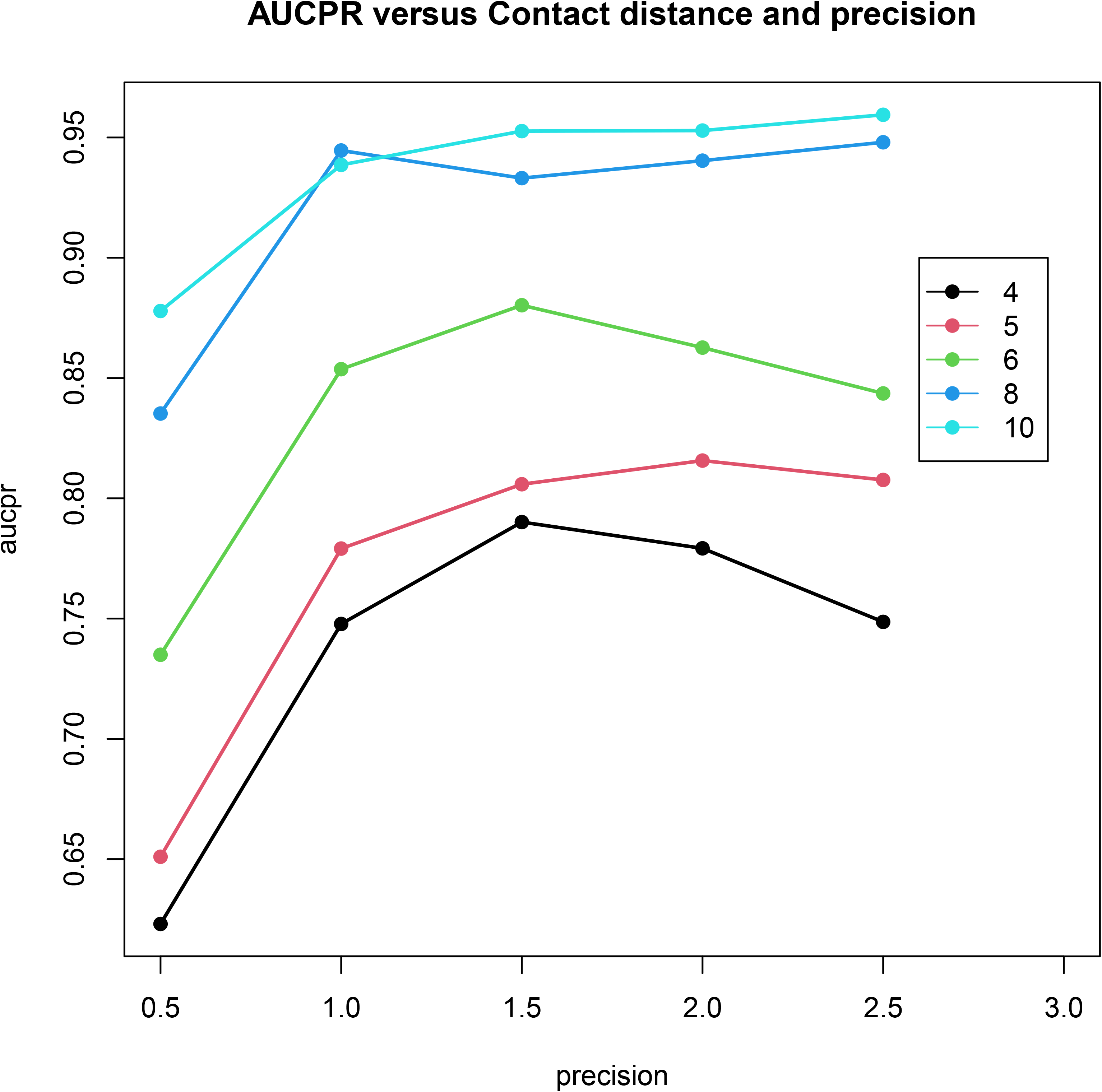

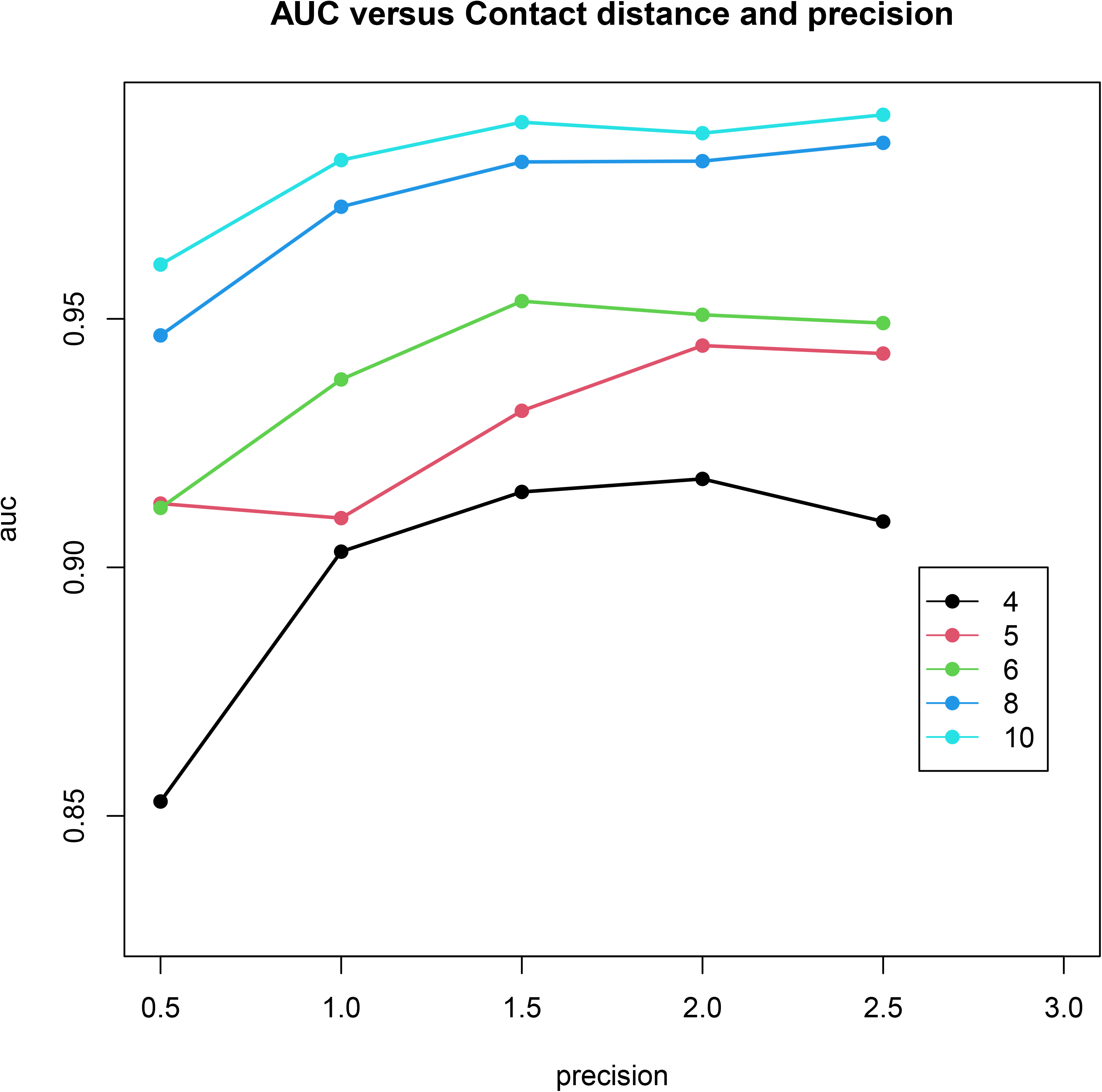

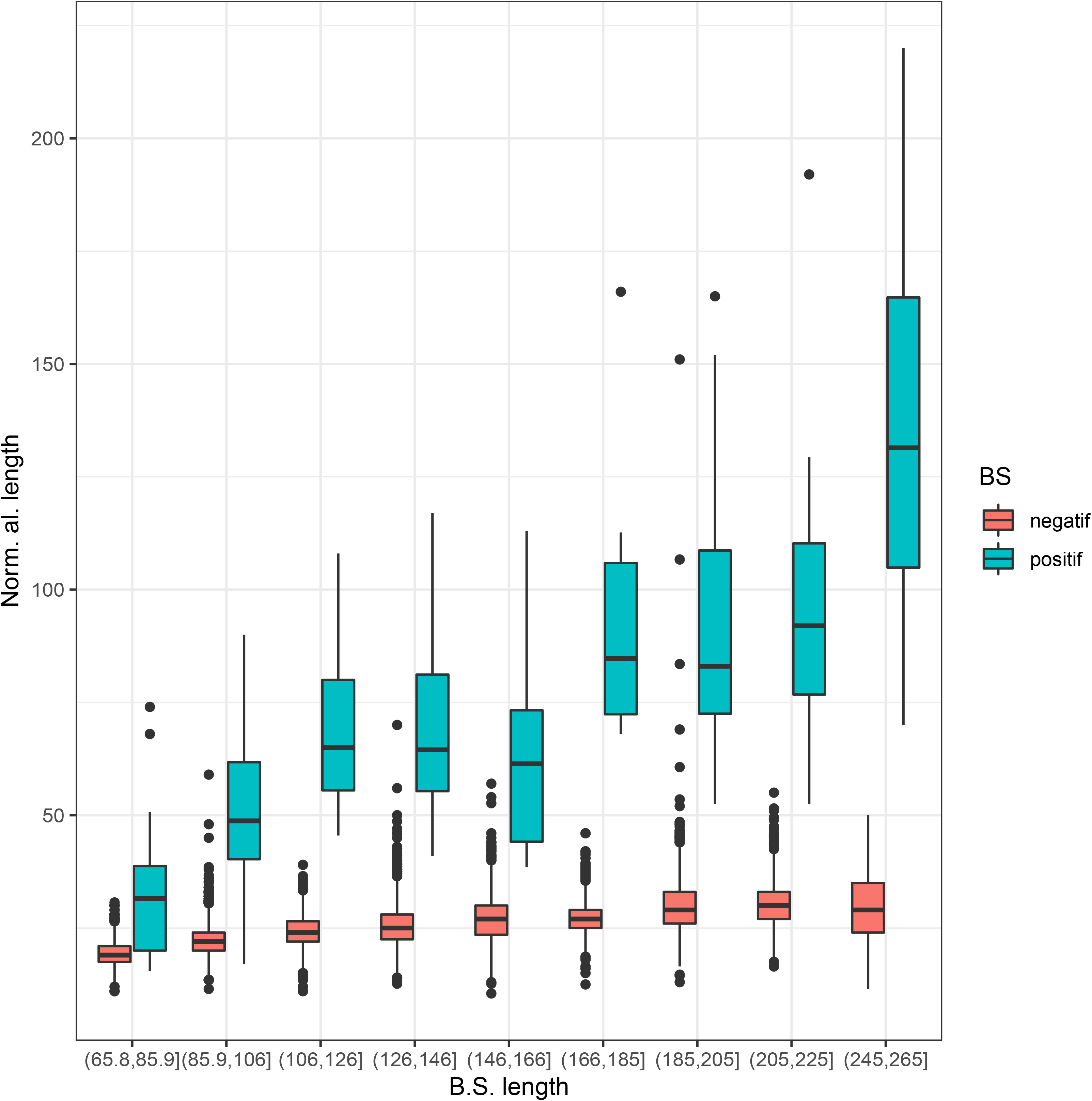

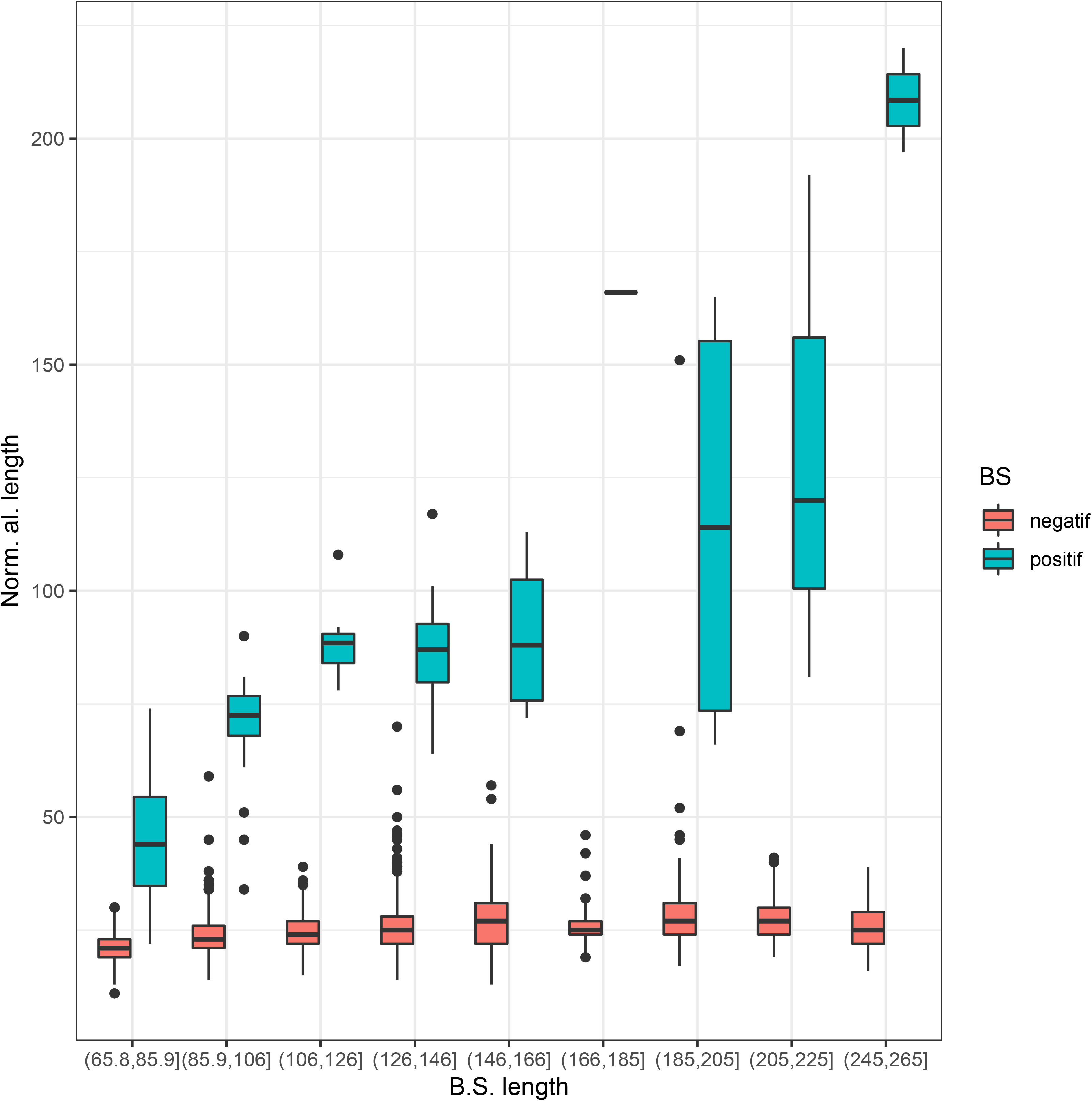

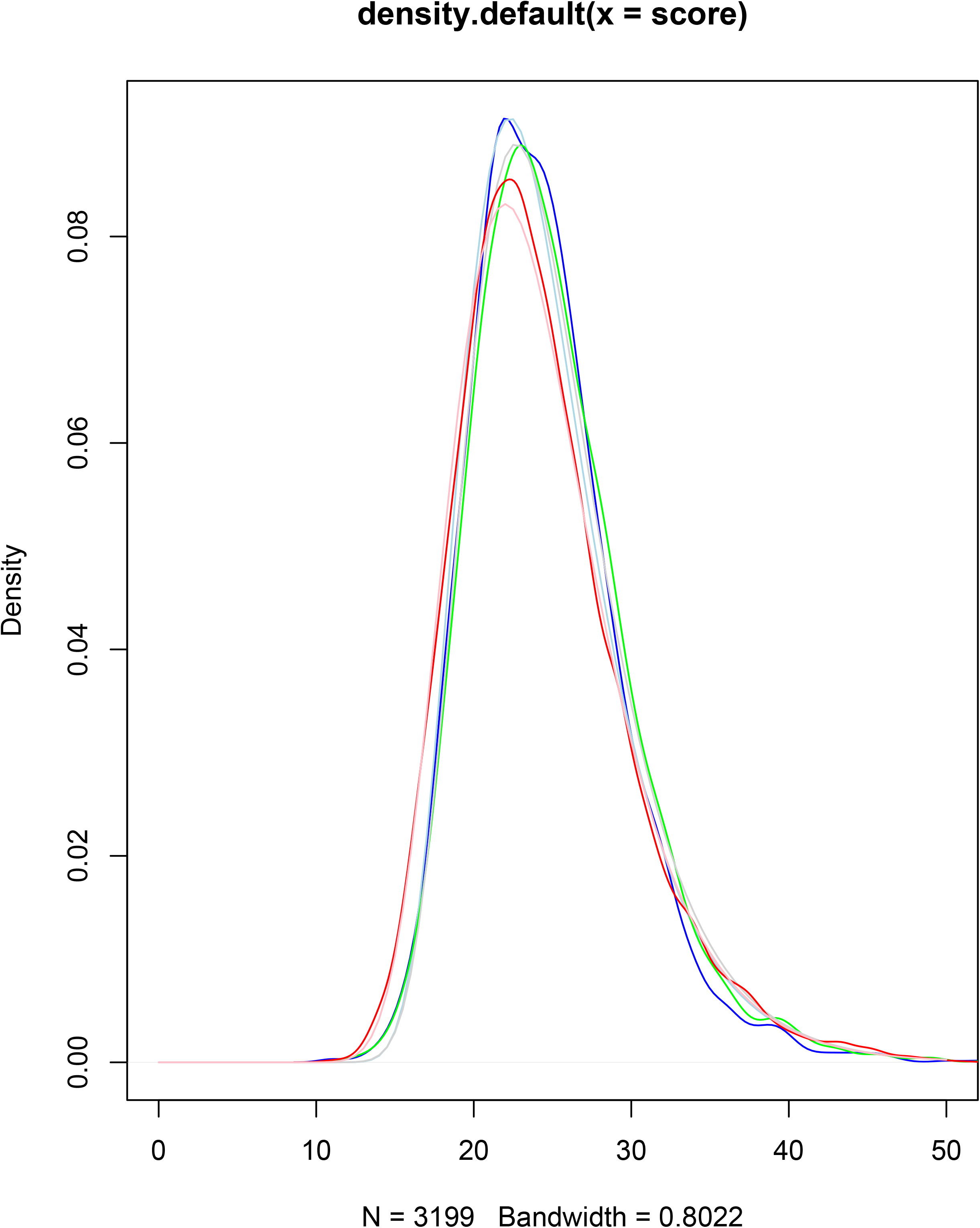

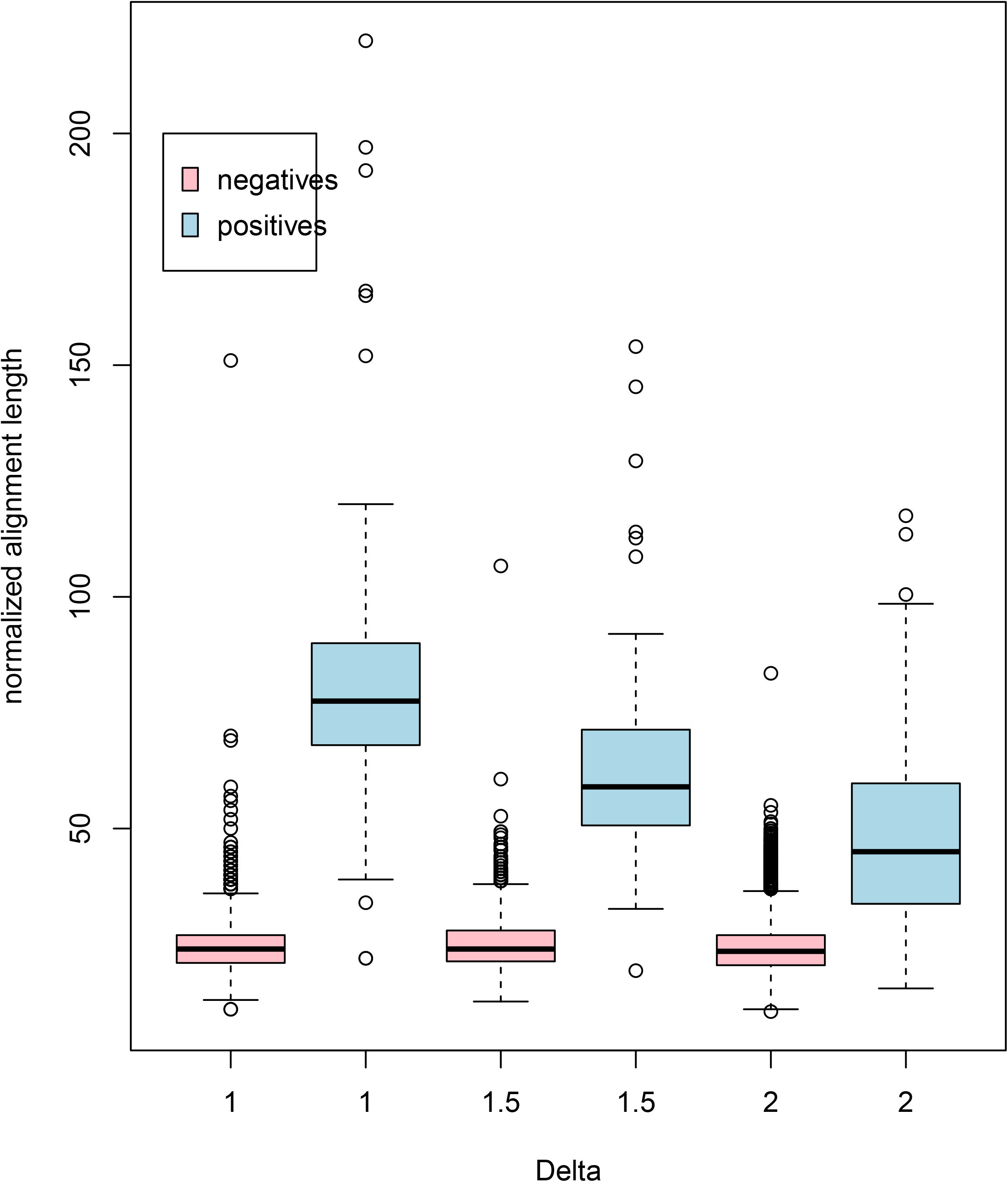

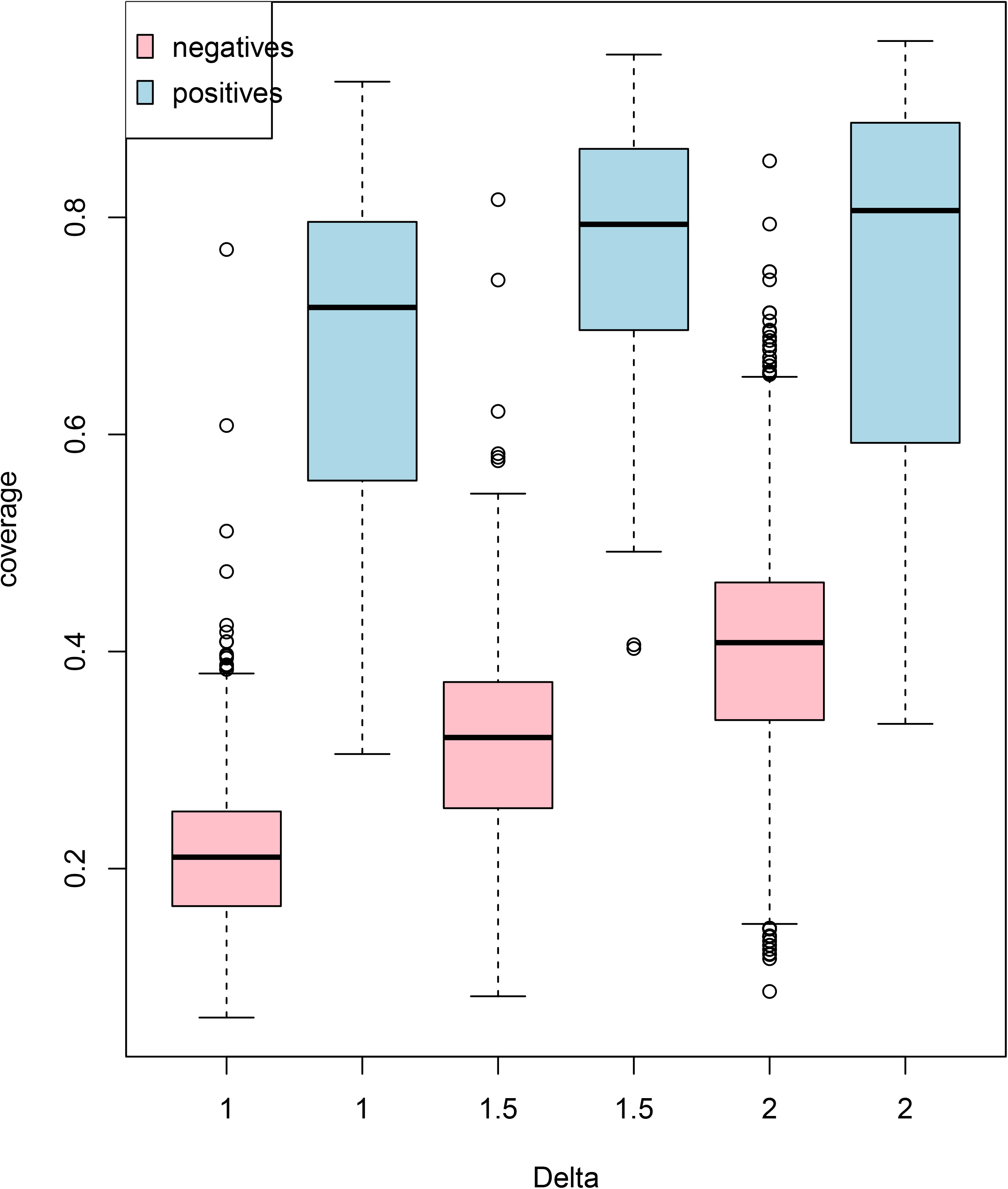

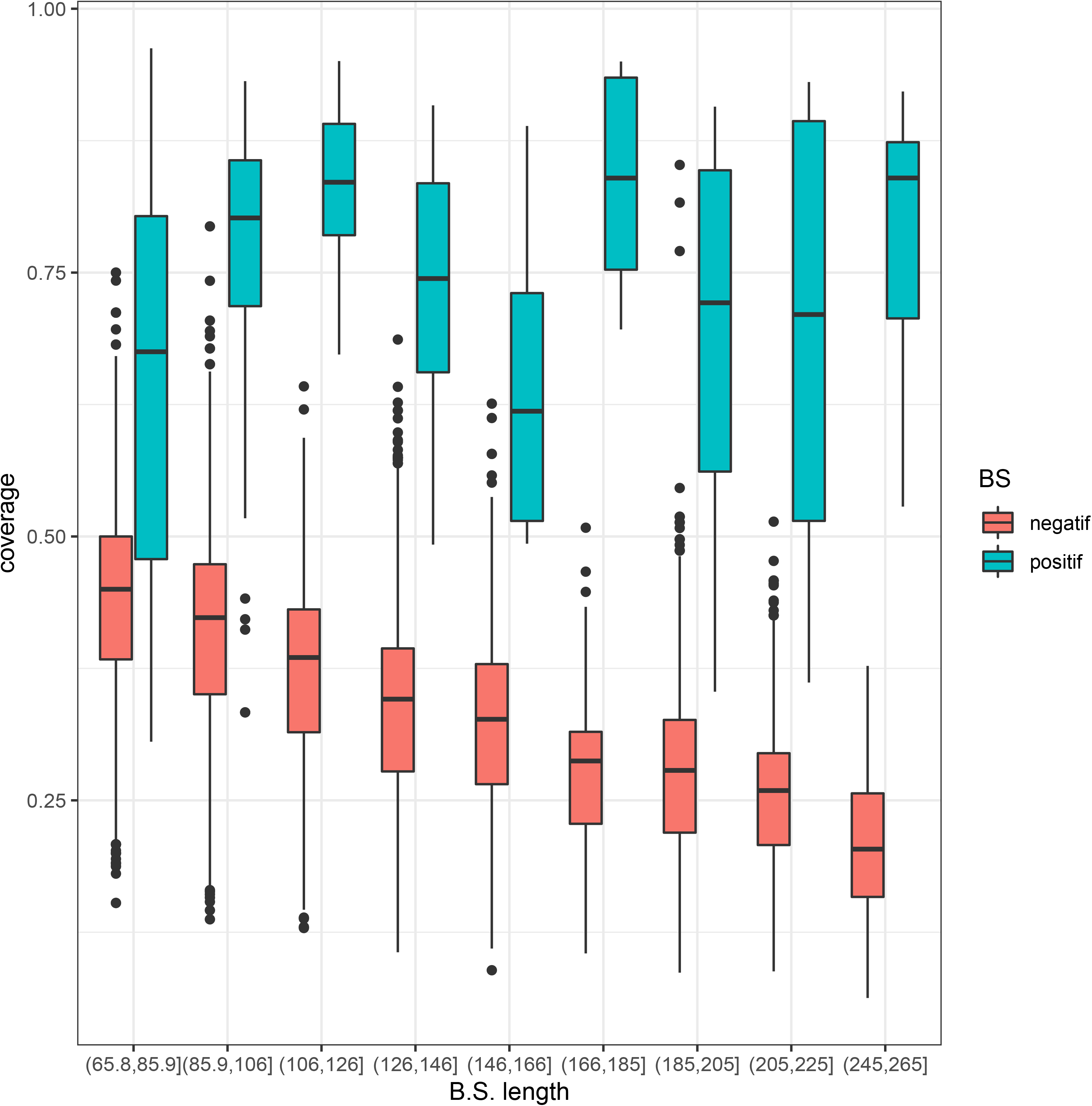

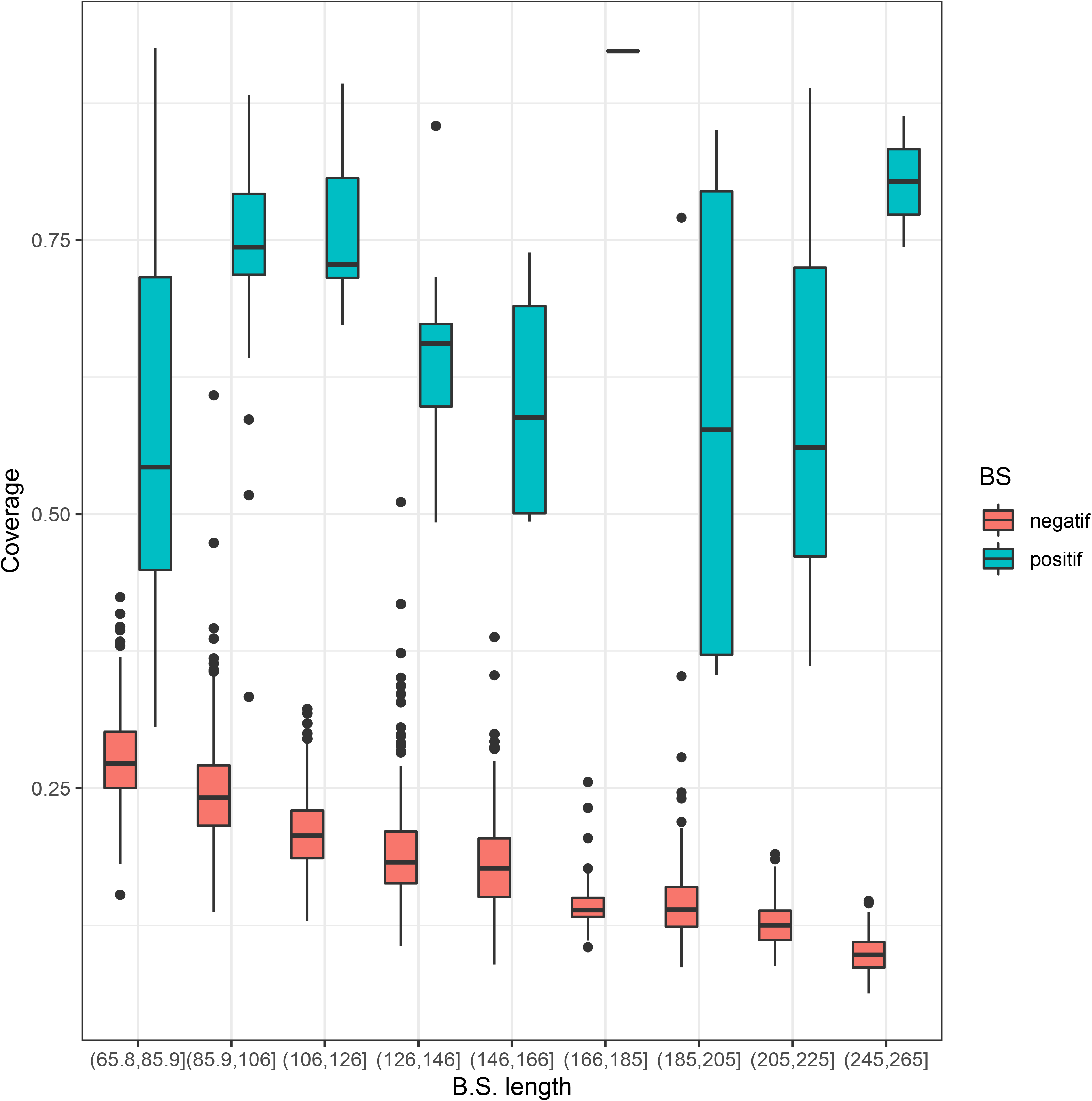

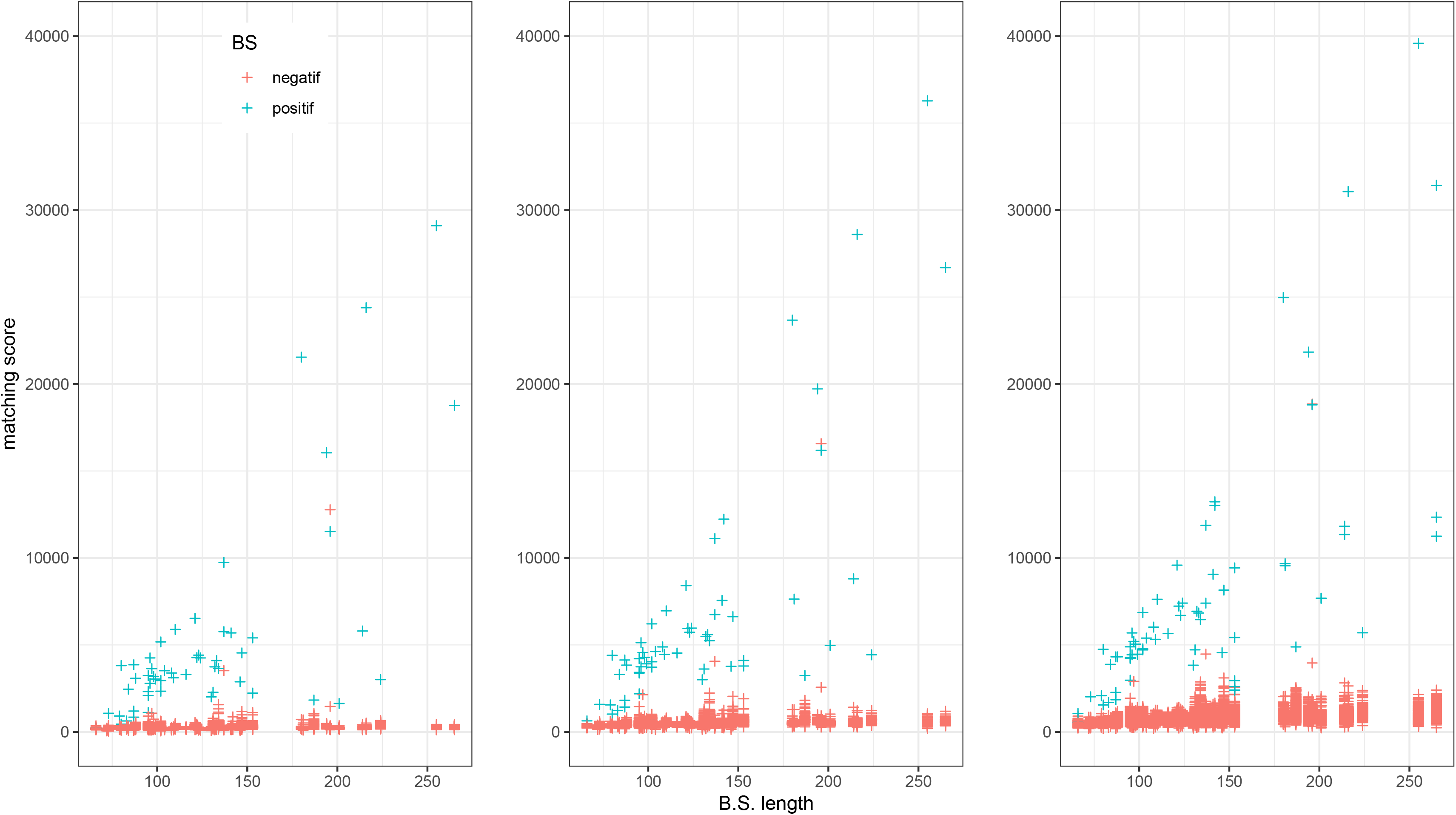

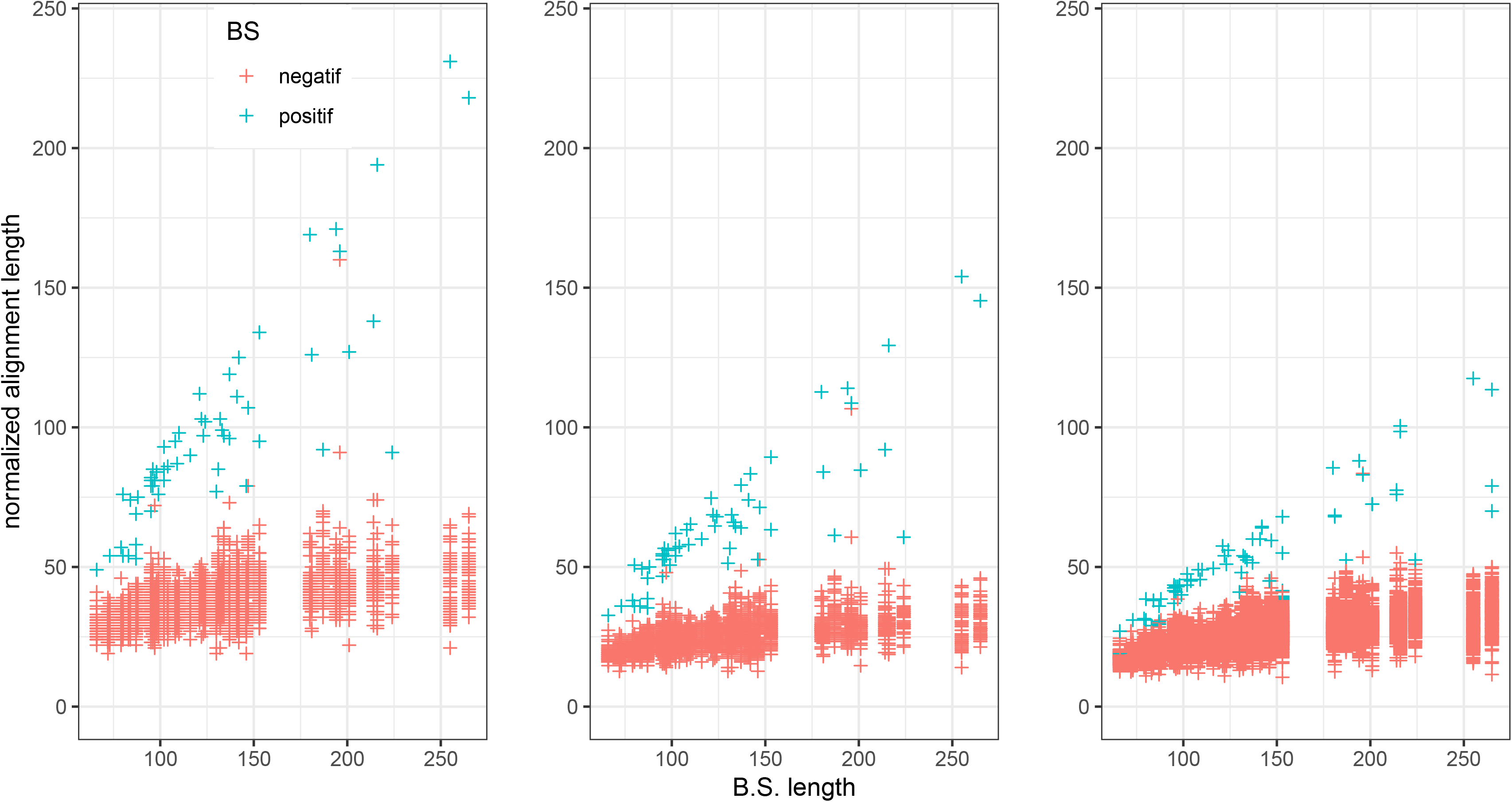

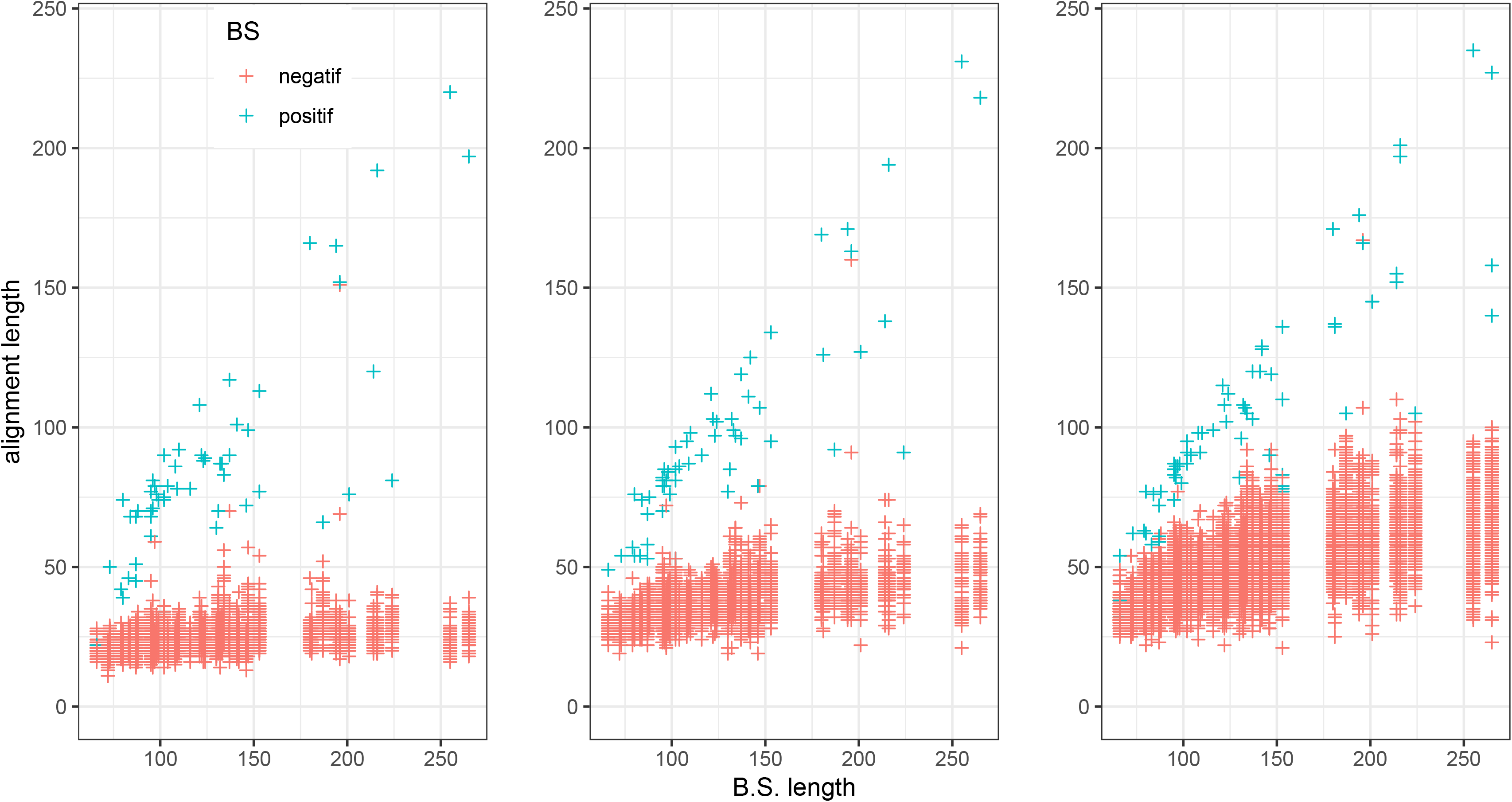

